# Actin-regulated Siglec-1 nanoclustering influences HIV-1 capture and virus-containing compartment formation in dendritic cells

**DOI:** 10.1101/2022.04.28.489919

**Authors:** Enric Gutiérrez-Martínez, Susana Benet, Nicolas Mateos, Itziar Erkizia, Jon Ander Nieto-Garai, Maier Lorizate, Carlo Manzo, Felix Campelo, Nuria Izquierdo-Useros, Javier Martinez-Picado, Maria F. Garcia-Parajo

## Abstract

The immunoglobulin-like lectin receptor CD169 (Siglec-1) mediates the capture of HIV-1 by activated dendritic cells (DC) through binding to sialylated ligands. These interactions result in a more efficient virus capture as compared to resting DCs, although the underlying mechanisms are poorly understood. Using a combination of super-resolution microscopy, single particle tracking and biochemical perturbations we studied the nanoscale organization of Siglec-1 on activated DCs and its impact on viral capture and its trafficking to a single viral-containing compartment. We found that activation of DCs leads to Siglec-1 basal nanoclustering at specific plasma membrane regions where receptor diffusion is constrained by Rho-ROCK activation and formin-dependent actin polymerization. Using liposomes with varying ganglioside concentrations, we further demonstrate that Siglec-1 nanoclustering enhances the receptor avidity to limiting concentrations of gangliosides carrying sialic ligands. Binding to either HIV-1 particles or ganglioside-bearing liposomes lead to enhanced Siglec-1 nanoclustering and global actin rearrangements characterized by a drop in RhoA activity, facilitating the final accumulation of viral particles in a single sac-like compartment. Overall, our work provides new insights on the role of the actin machinery of activated DCs in regulating the formation of basal Siglec-1 nanoclustering, being decisive for the capture and actin-dependent trafficking of HIV-1 into the virus-containing compartment.

## Introduction

Dendritic cells (DCs) are a specialized group of leukocytes that play an essential role in the innate and adaptive immunity through their function as antigen presenting cells (Banchereau and Steinman, 1998). DCs patrol peripheral tissues capturing invading pathogens, including viruses such as HIV-1, which are then degraded into antigens as the cells migrate towards the lymph nodes to activate T-cell responses. However, HIV-1 can also exploit DCs as vehicles to *trans*-infect CD4^+^ T cells, a process that allows the dissemination of the virus through cell-to-cell contacts (Cameron et al., 1992; Pope et al., 1994). Although immature DCs (iDC) express receptors and co-receptors required for HIV-1 infection (Granelli-Piperno et al., 1996; Turville et al., 2002), their infection rates in culture are lower than the ones for activated CD4^+^ T cells or macrophages (CAMERON et al., 1992; Granelli-Piperno et al., 1999, 1998; Pope et al., 1995). Moreover, iDCs direct part of the trapped viruses to the endolysosomal degradation pathway, which further decreases both the susceptibility of iDCs to infection and the load of viral particles that reach the cytoplasm to initiate their intracellular replication (Turville et al., 2004). As such, iDCs are inefficient *trans*-infecting T cells. In strong contrast, activation of DCs by lipopolysaccharide (LPS) or interferon (IFN), which are immunomodulatory signals present during HIV-1 infection, diminishes their susceptibility to infection but significantly increases the capture of HIV-1 and enhances the *trans*-infection of CD4^+^ T cells (Izquierdo-Useros et al., 2007; Wang et al., 2007).

The immunoglobulin-like lectin receptor CD169 (Siglec-1) is a key molecule in the HIV-1 capture and *trans*-infection of T cells by activated DCs (Izquierdo-Useros et al., 2012b; Puryear et al., 2013). Siglec-1 is a type I transmembrane protein that contains an amino-terminal V-set domain, which can directly bind to sialylated ligands like the ones present in gangliosides of the HIV-1 membrane (Crocker et al., 2007). In LPS-activated DCs (i.e., mature DCs, mDCs), once HIV-1 engages Siglec-1, a large amount of viral particles accumulate in a compartment connected to the extracellular milieu and enriched in several tetraspanin proteins (Garcia et al., 2005; Hyun et al., 2008; Izquierdo-Useros et al., 2012b, 2011; Puryear et al., 2013). The formation of this virus-containing compartment (VCC) reduces HIV-1 endolysosomal degradation while the accumulation of viruses enhances the viral load to *trans*-infect T cells (Izquierdo-Useros et al., 2012b).

Activation of DCs by LPS or IFNα upregulates the expression levels of Siglec-1, which partially explains their higher HIV-1 capture ability as compared to iDCs (Akiyama et al., 2015; Izquierdo-Useros et al., 2012b). However, multiple studies indicate that the spatial organization of plasma membrane receptors in immune cells as well as structural characteristics of their ligands also play an important role in the regulation of ligand/receptor interactions (Cambi et al., 2004; De Bakker et al., 2007; Gold and Reth, 2019; Neumann et al., 2019; Sherman et al., 2011; Torreno-Pina et al., 2016, 2014). For instance, the C-type lectin dendritic cell-specific intercellular adhesion molecule grabbing non-integrin (DC-SIGN), which also mediates the capture and internalization of several pathogens, including HIV-1 (Kwon et al., 2002), organizes in small nanoclusters in the plasma membrane of iDCs (Cambi et al., 2004; De Bakker et al., 2007). Such confined distribution increases the avidity of DC-SIGN for multimeric ligands by providing docking nano-platforms particularly efficient for the capture, internalization and degradation of viral particles (Cambi et al., 2004). Recently, another study established a link between the physical properties of synthetic glycopolymers carrying the mannose ligands for DC-SIGN and the internalization rates of the receptor [28]. Longer polymers with a higher number of mannose copies internalize faster than shorter ones, and polymer aggregation in the surface of particulate antigens diverge DC-SIGN receptors from endocytosis, directing them to invaginated pockets similar to the VCC harboring HIV-1 (Jarvis et al., 2019). In the context of Siglec-1, artificial liposomes carrying the ganglioside GM4, which contains a sialic acid moiety bound to a single galactose, are barely captured by mDCs (Izquierdo-Useros et al., 2012a). In contrast, liposomes with GM1, GM2, or GM3, whose sialic acid is bound to a lactose head group, efficiently bind to Siglec-1 with different concentration-dependent rates [29,30]. Overall, these findings indicate that the final outcome of viral capture and intracellular trafficking is the combined result of, on one hand, expression levels and nanoscale organization of the receptors and, on the other hand, concentration, spatial distribution and intrinsic structural properties of the ligands.

Here we used a combination of super-resolution stimulated emission depletion (STED) microscopy and single particle tracking (SPT) approaches to unravel the role of Siglec-1 nanoscale organization in the capture and trafficking of HIV-1 towards the VCC on activated DCs. Overall, our work reveals that distinctive components of the activated DC actin machinery regulate in a synergistic fashion the different stages of viral capture, trafficking and VCC formation, providing activated DCs with an increased capacity to *trans*-infect T cells.

## Results

### DC activation induces the formation of Siglec-1 nanoclusters with decreased mobility

The *trans*-infection capacity of DCs correlates with the expression levels of Siglec-1, which are increased upon cell activation with LPS or IFN (Izquierdo-Useros et al., 2012b; Puryear et al., 2013). However, the spatial organization of Siglec-1 receptors in resting or activated DCs is not yet known. To address whether DC maturation alters Siglec-1 distribution in the plasma membrane, we first examined the nanoscale organization of Siglec-1 by STED microscopy in iDCs and LPS-treated DCs (mDCs) differentiated from peripheral blood monocytes (PBMCs) (*Figure 1A and Figure 1-source data 1*). With a lateral resolution of ~ 80 nm (*figure supplement 1A*), we discriminated individual Siglec-1 fluorescent spots on the cell surface and measured their peak intensities. To quantify the number of Siglec-1 molecules per spot in the plasma membrane of iDC and mDC we relied on the intensity obtained from individual antibodies (Abs) on glass, corresponding to single molecules (*figure supplement 1B-D*) (Martínez-Muñoz et al., 2018). LPS-mediated DC maturation induced a higher fraction (~ 57% vs. ~37%) of Siglec-1 dimers and small nanoclusters with > 3 molecules/spot as compared to iDCs, where Siglec-1 was mainly found as monomers (~57% vs. ~22%) (*Figure 1A, B*). The increase in the average number of molecules/spot also coincided with an average increase in spot sizes (*Figure 1C*) and a significant increase in the proximity of Siglec-1 spots on mDCs as compared to iDCs (*Figure 1D*). To rule out that nanoclustering and their spatial proximity were simply a consequence of the increased expression of Siglec-1 upon DC maturation, we performed simulations of spots randomly distributed through the cell area using identical Siglec-1 molecular densities as obtained from the experimental data (*figure supplement 1E*). Interestingly, normalization of the experimental data to the randomized values preserved the significant differences between mDC and iDC in spot size, proximity between spots and number of molecules/spot (*figure supplement 1F-H and Supplementary figure S1-source data Fig. S1*). Thus, these results indicate the existence of real, active basal nanoclustering of Siglec-1 after DC maturation.

**Figure 1.**
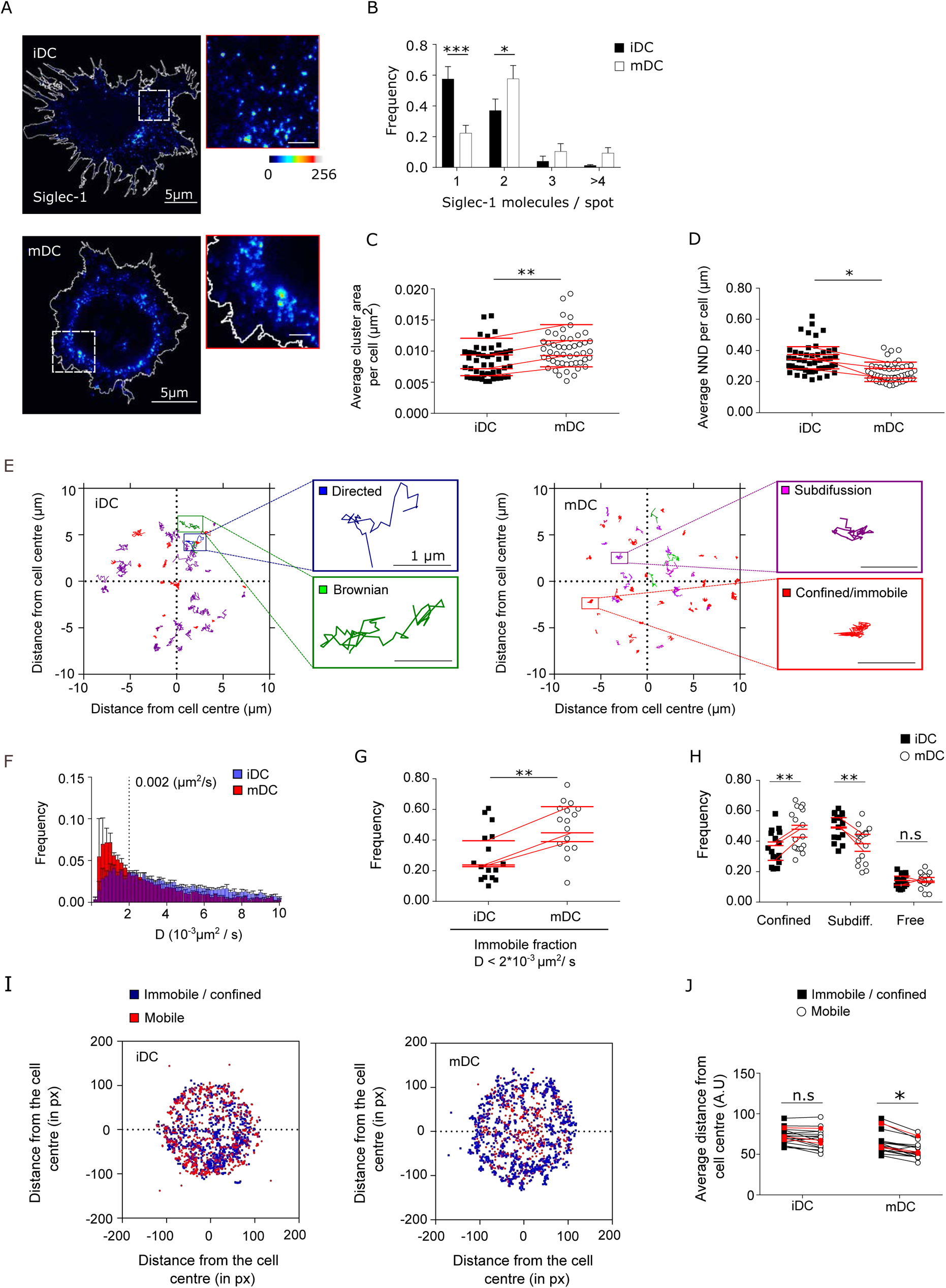
DC activation induces the formation of Siglec-1 nanoclusters with decreased mobility. (A) Representative STED images of Siglec-1 in iDCs and mDCs. The pseudo-color code denotes the intensity of individual Siglec-1 labeled spots, from monomer (dark blue) to nanocluster (red-to-white). The insets show enlarged regions highlighted by the white boxes on the main images (scale bar 1µm). (B) Frequency of the number of Siglec-1 molecules per spot, in iDC and mDC. Bars represent the mean ± S.E.M of 3 different donors (minimum of 9 cells/donor and condition). (C) Average size of Siglec-1 spots and (D) proximity between individual spots, calculated by measuring the nearest neighbor distance (NND) between spots per cell, in iDC and mDC. Each symbol in (C, D) corresponds to an individual cell, red lines are the average value on iDCs and mDCs for each donor (4 donors, 9 cells/donor and cell type). (E) Representative Siglec-1 trajectories as recorded on an iDC (left) and an mDC (right). The magnified insets show examples of different types of motion as classified by the MSS analysis. (F) Frequency of the diffusion coefficients for individual Siglec-1 trajectories on iDCs and mDCs. The dash vertical line corresponds to the diffusion threshold to separate immobile from mobile particles (0.002 µm^2^/s). Each data set represents the mean ± S.E.M of 3 donors (minimum of 3 cells and 83 trajectories/cell). (G) Fraction of immobile trajectories (< 0.002 μm^2^/s). Each symbol corresponds to an individual cell, red lines are the average value on iDCs and mDCs for each donor (3 donors). (H) Fraction of mobile trajectories (> 0.002 μm^2^/s) classified as confined, sub-diffusive or free. Each symbol corresponds to an individual cell, red lines show the average value on iDCs and mDCs for 3 donors analyzed. I) Plots showing the center of mass of individual Siglec-1 trajectories (average *x,y* position in all the frames of a given trajectory) in iDC and mDC. Blue dots correspond to immobile and confined trajectories, and red dots correspond to sub-diffusive and free trajectories. The graph shows all the trajectories analyzed for a minimum of 8 cells per condition from one donor. (J) Distance from the cell center of immobile/confined (black squares) and sub-diffusive/free (empty circles) trajectories in iDC and mDC. Black squares are paired to the empty circles within the same cell. Red symbols correspond to the average values of a minimum of 3 iDC and mDC cells per donor (3 donors).

The differences in Siglec-1 nanoscale organization between iDC and mDC might result from a difference in receptor mobility in the plasma membrane (Martínez-Muñoz et al., 2018; Torreno-Pina et al., 2016). To characterize the mobility of Siglec-1 we performed single particle tracking (SPT) of individual Siglec-1 molecules labeled at low density. Albeit different types of mobility were observed on both cell types (representative tracks in *Figure 1E*), DC maturation led to an overall reduction of Siglec-1 diffusion (*Figure 1F*) and increased the fraction of immobile receptors (diffusion values < 0.002 μm^2^/s), as compared to iDCs (~28% vs ~48%) (*Figure 1G*). We further used the momentum scaling spectrum (MSS) analysis to classify the type of motion of the mobile trajectories (Ferrari et al., 2001; Sbalzarini and Koumoutsakos, 2005). We found a significant increase in the fraction of confined trajectories on mDCs as compared to iDC (~47% vs. ~35%), and no differences in the fraction of free mobile trajectories between both cell types (*Figure 1H*). Moreover, immobile and/or confined trajectories in mDC were preferentially located close to the cell edges (*Figure 1I and J*) as compared to the more even distribution observed in iDC, suggesting that upon DC activation Siglec-1 diffusion becomes spatially constrained in specific regions of the plasma membrane. Altogether, these results indicate that LPS activation of DCs not only increases Siglec-1 expression, but importantly, it also modulates its nanoclustering, lateral mobility and overall spatial distribution on the plasma membrane.

### Formin-dependent actin polymerization regulates Siglec-1 nanoscale organization and mobility

Previous studies have demonstrated an essential role for the actin cytoskeleton modulating the capture of HIV-1 and the formation of the VCC by mDCs (Izquierdo-Useros et al., 2011; Ménager and Littman, 2016). Although Siglec-1 does not have known intracellular motifs of actin interaction (Bornhöfft et al., 2018), it has been documented that the actin cytoskeleton can regulate the nanoscale organization of different membrane receptors, and even lipids, that do not directly bind to actin [36–39]. To assess the role of actin in the spatial organization of Siglec-1, we performed dual color STED imaging of Siglec-1 together with actin in iDCs and mDCs (*Figure 2A and Figure 2-source data 2*). When we traced the intensity distributions of actin and Siglec-1 from the cell center we observed a good spatial correlation of both markers in iDC and mDC (*Figure 2B*). However, consistent with our previous analysis showing the preferential distribution of confined Siglec-1 molecules close to the cell edges, Siglec-1 and actin were highly enriched near the edges of mDC as compared to iDC, suggesting a role for actin in the regulation of Siglec-1 nanoclustering and its spatial confinement.

**Figure 2.**
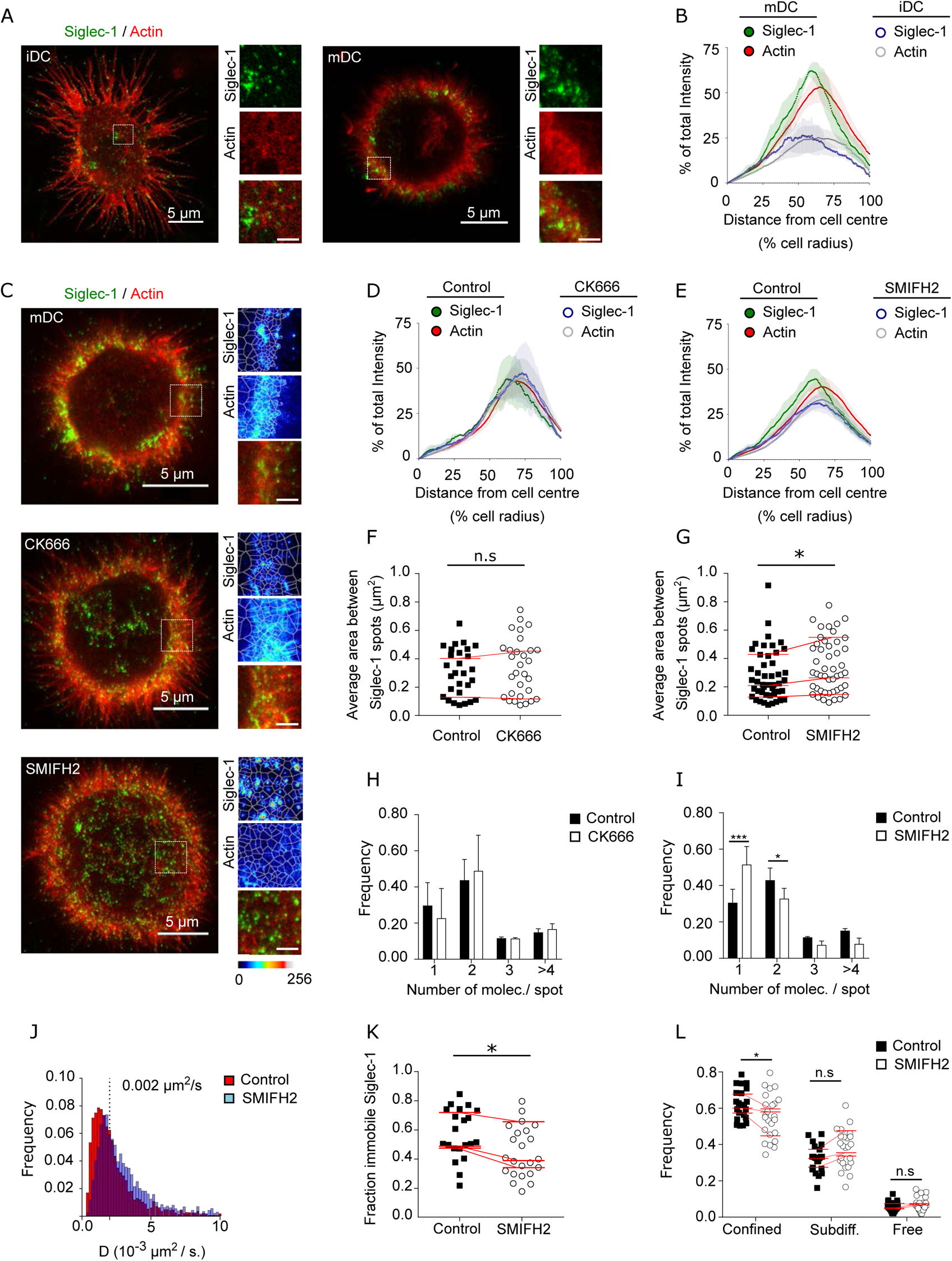
Formin-dependent actin polymerization regulates Siglec-1 nanoscale organization and mobility. (A) Representative dual color STED images of iDC (left) and mDC (right) stained with Siglec-1 and actin (labeled with SiR-actin). Insets show magnified areas with pseudo-color-coded intensities of Siglec-1 (upper), actin (middle) and merged image (lower) (scale bar 1µm). (B) Siglec-1 and actin intensity from the cell center towards the edge expressed as percentage of total intensity for each marker in iDC and mDC. Lines represent the mean ± S.E.M of 3 different donors (minimum of 9 cells/donor and condition). C) Representative STED images of mDC stained for Siglec-1 and actin, together with enlarged insets, in control conditions and after 1 h treatment with CK666 (100 µM) or SMIFH2 (25 µM). The white lines in the insets delimit the Voronoi areas between adjacent Siglec-1 spots; the smaller the areas, the closer single Siglec-1 spots reside from each other (Scale bars insets: 1 µm). (D-E) Siglec-1 and actin intensity from the cell center towards the edge expressed as percentage of total intensity for each marker in control mDC and treated with CK666 (D) or SMIFH2 (E). Lines represent the mean ± S.E.M of 2 (D) and 4 (E) different donors (minimum 10 cells/donor and condition). (F) Average Voronoi areas between contiguous Siglec-1 spots per cell in control and CK666 treated cells (2 donors, at least 10 cells/donor). Red lines connect the average of all the cells in each donor. (G) Similar to (F) for control and SMIFH2 treated cells (3 donors, at least 10 cells/donor). (H) Frequency histogram of the number of Siglec-1 molecules/spot in control and CK666-treated mDC. Mean ± S.E.M. of a minimum of 10 cells/donor from 2 donors. (I) Similar to (H) for control and SMIFH2 treated cells (3 donors, at least10 cells/donor). (J) Frequency of Siglec-1 diffusion coefficients in control mDC (n=8 cells, 4424 trajectories) and cells treated with SMIFH (n=8 cells, 5002 trajectories) from one donor. (K) Fraction of immobile trajectories in control and SMIFH2 treated mDCs. A total of 21 (control) and 24 (SMIFH2) cells were quantified from 3 donors (minimum 6 cells and 1261 trajectories/donor and condition). Red lines connect the average of each individual donor. (L) Fraction of mobile trajectories classified as confined, sub-diffusive or free in control and SMIFH2-treated mDC. Each symbol corresponds to an individual cell, red lines show the average value of all cells from 3 donors (minimum 6 cells/ donor and condition).

To further test this hypothesis, we first treated mDCs with Cytochalasin D (CytoD), a drug that inhibits actin polymerization. In these conditions, Siglec-1 staining dispersed throughout the plasma membrane (*figure supplement 2A and B and Supplementary figure S2-source data Fig. S2*) and the fraction of Siglec-1 receptors found in dimers or small nanoclusters decreased in favor of a significant increase of monomers (*figure supplement 2C*). Because cortical actin mainly depends on two different polymerization mechanisms, one Arp2/3-dependent which forms branched actin, and another dependent on formins which produce filamentous actin bundles (Rottner et al., 2017), we selectively inhibited each of these mechanisms using CK666 (Arp2/3 inhibitor) or SMIFH2 (formin inhibitor) in mDCs (*Figure 2C*). The association of Siglec-1 high density regions with cortical actin near the cell edges was not affected by Arp2/3 inhibition, but decreased when we inhibited formins (*Figure 2D, E*). Such dispersion was further quantified using a tessellation-based analysis to define areas between adjacent Siglec-1 spots (the smaller the areas, the higher the density). Whereas Arp2/3 inhibition did not affect the proximity between Siglec-1 spots (*Figure 2F*), formin inhibition significantly increased the separation between individual spots consistent with their spatial dispersion (*Figure 2G*). Importantly, formin inhibition, but not Arp2/3 inhibition, reduced the basal Siglec-1 nanoclustering as compared to control conditions (*Figure 2H and I*).

These results indicate that Siglec-1 nanoclustering in activated DCs relies on a formin-dependent actin polymerization mechanism of confinement. These findings were further substantiated by SPT analysis. Although the mobile fraction of Siglec-1 (above 0.002 µm^2^ /s) did not show significant differences in the diffusion coefficients (*Figure 2J*), we observed a significant decrease in the fraction of immobile and confined trajectories upon formin inhibition (*Figure 2K and L*). Moreover, consistent with the dissociation of Siglec-1 from the regions enriched in cortical actin, formin inhibition also led to a more even distribution of immobile and mobile Siglec-1 receptors through the cell area (*figure supplement 2D and E*).

To further demonstrate the relationship between formins and Siglec-1 nanoscale organization, we measured Siglec-1 distribution in a physiological situation in which different types of actin pools simultaneously occur at specific regions of the plasma membrane. Activated DCs in confined environments, when subjected to chemokine stimuli, migrate in a process that requires actin polarization at the leading edge (mainly dependent on Arp2/3 branched actin) and at the cell rear (formin-dependent filamentous actin) (Jolly et al., 2007; Lämmermann et al., 2009; Paluch et al., 2016; Vargas et al., 2016). We thus squeezed activated DCs between a layer of agarose and a coverslip with a homogeneous concentration of the chemokine CCL19, and subsequently fixed and stained the cells for Siglec-1 and actin. The spatial constrain of mDCs in this semi-3D environment was optimal to observe in the same plane a clear polarization of a protrusive lamellipodium and a contractile trailing edge (*figure supplement 2F*). Whereas actin staining was isotropic at the front and cell rear (*figure supplement 2G*), Siglec-1 was remarkably enriched at the formin-dependent trailing edge (*figure supplement 2H*). Moreover, a significant enhancement of Siglec-1 nanoclusters was detected at the trailing edge as compared to the lamellipodium at the front (*figure supplement 2I*). Collectively, these results indicate that formin-dependent cortical actin regulates the nanoscale organization, membrane lateral confinement and diffusion of Siglec-1 in activated DCs.

### Siglec-1 confinement and nanoclustering occurs in polarized regions of the plasma membrane characterized by RhoA activity

The previous results suggest differences in the organization of the actin cytoskeleton between mDCs and iDCs, which might impact on the overall spatiotemporal organization of Siglec-1. Recent studies have shown that DC maturation increases Rho activity in polarized regions of the membrane, which in turn enhances formin-dependent actin polymerization in filamentous bundles (Vargas et al., 2016). Thus, to assess the existence of a basal polarization of different actin polymerization mechanisms regulating the confinement of Siglec-1, we first imaged the distribution of Siglec-1 together with actin and the phosphorylated form of ezrin-radixin-moesin (pERM), a well-known actin-associated marker that becomes activated downstream of RhoA (*Figure 3A and Figure 3-source data 3*). As previously reported in LPS-activated macrophages (Di Pietro et al., 2017), we detected an increase in the levels of pERM in mDCs as compared to iDCs (*figure supplement 3A and Supplementary figure S3-source data Fig. S3*). Most importantly, we observed a clear polarization of Siglec-1 to cell regions closer to the basal membrane in mDCs (i.e., within the first 3-5 μm) (*Figure 3B*), and characterized by the abundance of filopodia enriched in pERM (*Figure 3C and D*).

**Figure 3.**
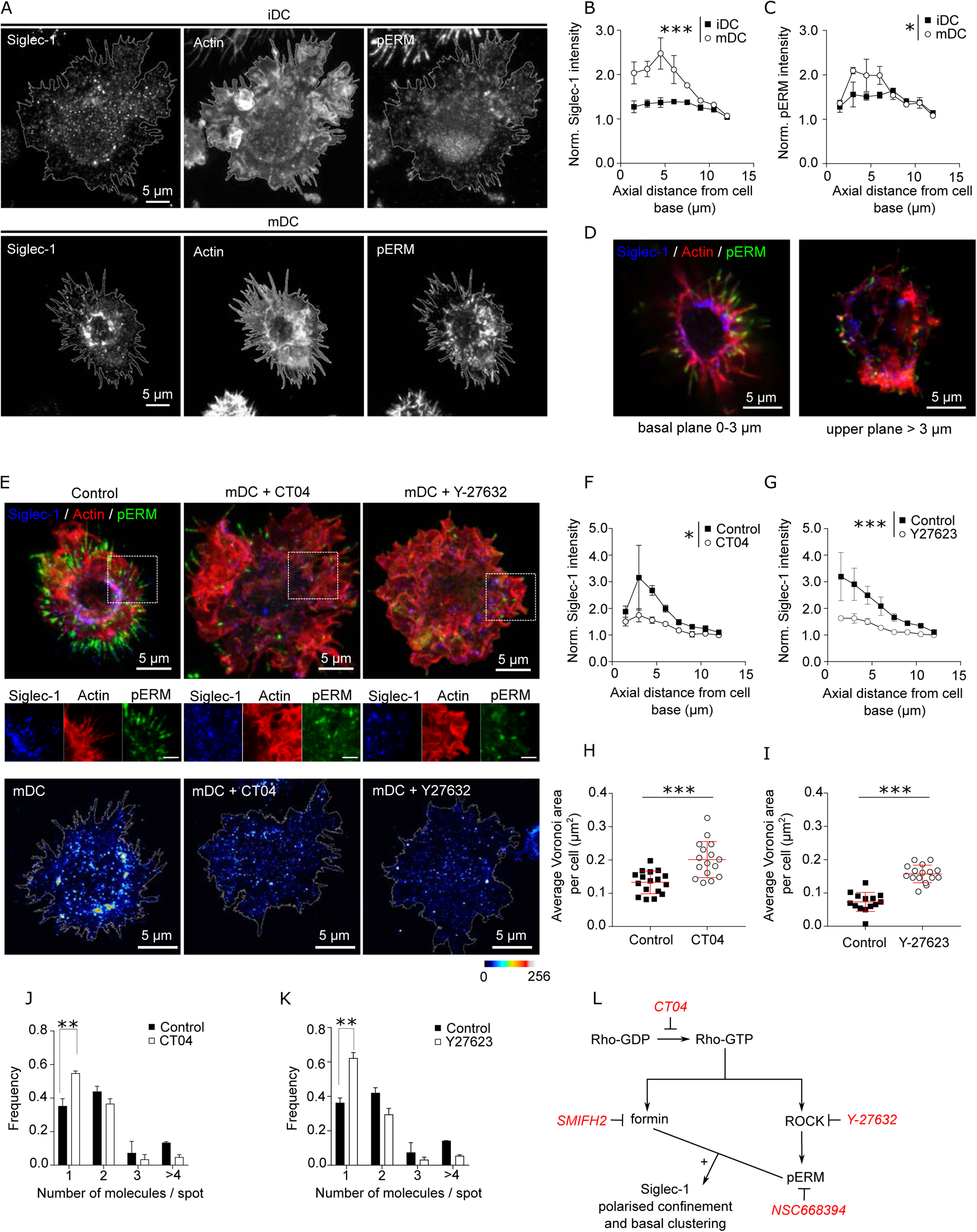
Siglec-1 confinement and basal nanoclustering occurs in polarised regions of the plasma membrane characterized by RhoA activity. (A) Representative confocal maximum intensity projections images of iDC (upper row) and mDC (lower row) immunostained with anti-Siglec-1, SiR-actin and pERM. (B-C) Plots of the axial distribution from the cell base of Siglec-1 (B) and pERM (C) intensities. Values in each plane show the mean intensity in steps of 1.5 µm from the cell base, normalized by the minimum mean intensity within the whole stack. Each symbol represents the mean ± S.E.M from two donors (at least 10 cells per condition). (D) Confocal projections of the basal (0-5 µm) and the apical (>5 µm) parts of the mDC example shown in (A). (E) Top panels: confocal maximum intensity projections images of control mDC and cells treated for 3 h with CT04 (2 µg/ml), or for 1h with Y-27632 (30 µM), fixed and immunostained with anti-Siglec-1, SiR-actin and pERM. Middle panels: enlarged images from the highlighted regions of the top images (scale bar: 2 µm). Bottom panels: STED images of mDCs treated as described above and immunostained with anti-Siglec-1. (F-G) Siglec-1 intensity as function of axial distance from the cell base in control mDC and cells treated with CT04 (F) and Y-27623 (G). Results show the mean ± S.E.M of two donors, with at least 7 (F) and 9 (G) cells/ donor and condition. (H-I) Average voronoi areas between contiguous Siglec-1 spots in control mDC, and cells treated with CT04 (H) and Y-27623 (I). Each dot shows the average Voronoi area per cell; results show the average ± S.D. of one representative experiment (n=2) with a minimum of 16 (H) and 14 (I) cells. (J-K) Number of Siglec-1 molecules/spot in control mDCs and cells treated with CT04 (J) and Y-27623 (K). Mean ± S.E.M of 2 donors, with a minimum of 10 (J) and 9 (K) cells/donor. (L) Scheme of the different pathways downstream of Rho activation targeted by different inhibitors. Statistics in the legends of panels B, C, F, G correspond to the significance of a Two-way ANOVA test depending on maturation status (iDC vs mDC in A,B) or treatment (control vs. CT04 or Y-27623 in F, G).

Next, we assessed if Rho activity was involved in the polarized distribution of Siglec-1 in mDCs. Pharmacological inhibition of RhoA (CT04) as well as of its downstream effector, the Rho-associated protein kinase ROCK (Y-27632), decreased the levels of pERM in mDCs (*figure supplement 3 and, C*) and importantly, disrupted the polarized confinement of Siglec-1 in mDCs, resulting in a more even distribution throughout the plasma membrane (*Figure 3E-G*). Moreover, RhoA and ROCK inhibition led to a significant decrease in the density of Siglec-1 nanoclusters, as determined by STED (*Figure 3E*, bottom panels; and *Figure 3H and I*), and to a reduced basal nanoclustering (*Figure 3J and K*), with values similar to those measured in iDCs. Significantly, treatment of iDCs with a RhoA activator (CN03), led to an increase of pERM levels (*figure supplement 3D and E*), Siglec-1 and pERM polarization in the basal plane of the cells (*figure supplement 3F and G*) and induction of basal Siglec-1 nanoclustering (*figure supplement 3H*) fully mirroring the results obtained on activated DCs. Together, these data strongly indicate that basal nanoclustering of Siglec-1 on activated DCs is the consequence of higher basal levels of RhoA activation, as compared to resting DCs.

It has been reported that the spatiotemporal organization of several receptors is influenced by the presence of additional membrane receptors that work as pickets in association with pERM (Freeman et al., 2019, 2018; Kalay et al., 2014; Kusumi et al., 2012; Trimble and Grinstein, 2015). Given our results, we thus checked the effects of ezrin inhibition in the spatial organization of Siglec-1 on mDCs. Pharmacological inhibition of ERM (NSC668394) led to a significant drop in the levels of pERM (*figure supplement 3I and J*), significantly disrupted the spatial polarization of Siglec-1 (*figure supplement 3K*) and reduced Siglec-1 nanoclustering (*figure supplement 3L*). Notably, formin inhibition did not alter the basal levels of pERM (*figure supplement 3M*) but disrupted the polarized distribution of Siglec-1 (*figure supplement 3N*). Hence, our results strongly indicate that Rho activity in mDCs regulates Siglec-1 basal nanoclustering and its spatial confinement in polarized regions of the plasma membrane through two necessary, but independent mechanisms: formin activation and ROCK-mediated ERM phosphorylation (*Figure 3L*).

### Siglec-1 basal nanoclustering increases the capture of HIV-1 particles and the avidity to gangliosides carrying sialic acid ligands

Previous reports have highlighted the importance of nanoscale organization of viral receptors on the capture of viral particles by providing multimolecular docking platforms with high avidity for viral ligands (Cambi et al., 2004). To assess the relevance of Siglec-1 nanoscale organization in the capture of HIV-1, we pulsed control, SMIFH2 or CT04-treated mDCs for 5 min with viral-like particles (VLPs). Inhibition of formin-mediated actin polymerization (*Figure 4A and B and Figure 4-source data 4*) or Rho activity (*figure supplement 4A and B and Supplementary figure S4-source data Fig. S4*), respectively, which disperse Siglec-1 nanoclustering and its spatial distribution, significantly decreased the immediate binding of VLPs to Siglec-1. Confocal stacks taken from cells immunostained with Siglec-1 without permeabilization revealed no significant differences in the total plasma membrane levels of Siglec-1 in control, SMIFH2 or CT04 treated cells (*Figure 4C* and *figure supplement 4C*), indicating that receptor availability was not responsible for the decrease in VLP capture. Remarkably, when we analyzed the nanoscale organization of Siglec-1 on 5 min-VLP-pulsed cells, we found that Siglec-1 nanoclusters colocalizing with VLPs had significantly higher number of molecules/spot as compared to non-colocalizing Siglec-1 nanoclusters, in both control and SMIFH2 treated cells (*Figure 4D*). These results suggest that both, formin and Rho-dependent Siglec-1 nanoclustering, facilitate the capture of VLPs by providing high-avidity docking platforms to viral ligands.

**Figure 4.**
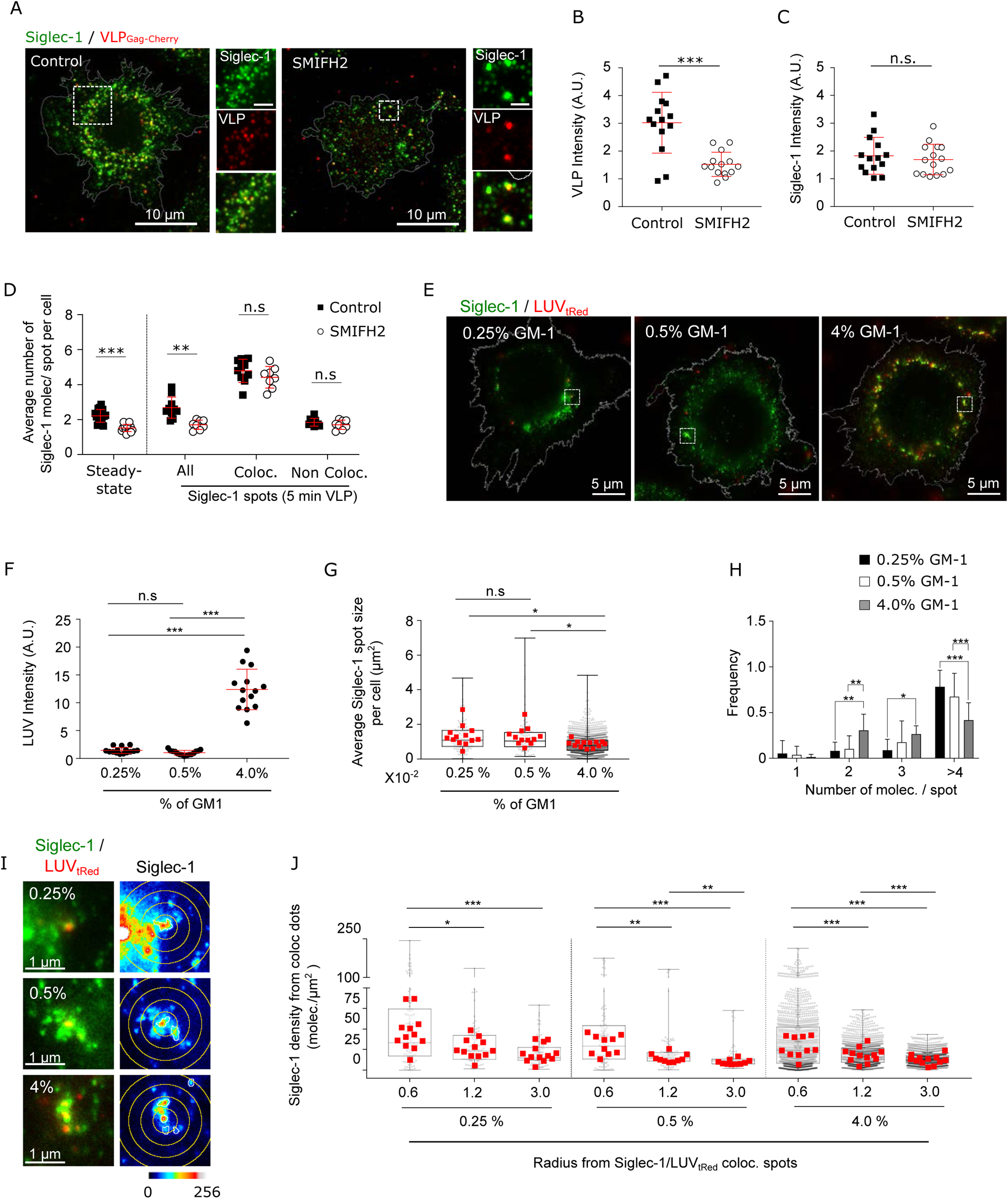
Siglec-1 basal nanoclustering increases the capture of HIV-1 particles and the avidity to gangliosides carrying sialic acid ligands. (A) Confocal images of control mDC, and cells treated with SMIFH2, pulsed for 5 min with VLP-Gag-Cherry and immunostained with anti-Siglec-1. Insets on the right correspond to enlarged views of the regions highlighted in the main images (scale bars: 2 µm). (B) VLP Intensity in control and SMIFH2 treated cells, after 5 min of VLP capture. Mean ± S.D. of one representative experiment (n=2) with 14 cells per condition. (C) Levels of Siglec-1 in control and SMIFH2 treated cells, after 5 min of VLP capture. Mean ± S.D. of one representative experiment (n=2), 14 cells per condition. (D) Average number of Siglec-1 molecules/spot per cell in control and SMIFH2 treated mDC, at steady-state and after 5 min of VLP capture. Siglec-1 data after 5 min VLP exposure are further separated according to whether Siglec-1 spots colocalize (or not) with VLP particles. Data correspond to the mean + S.D of one representative experiment (n = 3 for the absence of VLP and n = 2 for the 5 minutes VLP) in which a minimum of 8 cells per experiment and condition were analyzed). For each experiment a minimum of 8 cells per condition was (E) Representative dual color STED images of mDC immunostained with anti-Siglec-1 after 5 min pulse with tRed-LUVs carrying different GM-1 concentrations. Squares are shown as magnified regions in panel (I). (F) LUV intensity in mDC after 5 min of LUV capture. Mean ± S.D. of a minimum of 14 cells per condition. (G) Sizes of Siglec-1 spots colocalizing with LUVs. Small dots represent each spot analyzed and red squares correspond to the average size of all colocalizing spots in a cell (at least 12 cells per condition). (H) Number of Siglec-1 molecules per spot colocalizing with LUVs. Results correspond to the mean ± S.D. of a minimum of 12 cells per condition. (I) Magnified STED images of Siglec-1 colocalizing with LUVs carrying different GM-1 concentrations. Circles with different radii on the Siglec-1 images are drawn starting from the center of each colocalizing spot. (J) Siglec-1 density (i.e., number of Siglec-1 molecules/µm^2^) at different radius from Siglec-1 spots colocalizing with 0.25% to 4% GM-1 LUVs (each small dot is an individual colocalizing spot, red squares are the mean values in each cell analyzed, 12 cells per condition).

To further substantiate this hypothesis, we generated artificial Large Unilamellar Vesicles (LUVs) (~150nm in diameter) mimicking the lipid composition of the HIV-1 envelope, but with a gradient of concentrations of the Siglec-1 sialic-acid containing ligand GM-1 (from 0.25% to 4%). We pulsed mDCs for 5 min with LUVs and recorded dual color STED images of Siglec-1 and LUVs labeled with texasRed (tRed) (*Figure 4E*). As expected, the capture of LUVs was strongly dependent on the GM-1 concentration, being highest at 4% GM-1 (*Figure 4F*). Remarkably, LUVs with low levels of GM-1 (0.25% and 0.5%) showed a significant preference for binding to larger Siglec-1 nanoclusters and with higher number of molecules per spot, as compared to the LUVs with high GM-1 levels (4%) (*Figure 4G and H*). These data indicate that Siglec-1 clustering contributes to overcome the decrease in ligand availability. Moreover, LUVs preferentially bound to regions with high density of Siglec-1 receptors, an effect that was more pronounced for LUVs carrying low GM-1 levels (*Figure 4I and J*). As a whole, these results indicate that basal Siglec-1 nanoclustering and aggregation in highly dense regions of the membrane constitutes a physical mechanism to increase HIV-1 capture efficiency by increasing the avidity of Siglec-1 receptors to gangliosides carrying sialic acid ligands.

### Interaction with HIV-1 particles induces Siglec-1 changes at the nano- and meso-scale towards the formation of the VCC

It is known that initial virus binding in mDCs is followed by a progressive polarized movement of the viruses towards the VCC (Hyun et al., 2008; Izquierdo-Useros et al., 2011). To enquire if iDC and mDC show differences in the events that occur after initial binding of the virus to Siglec-1, we visualized and compared the spatial evolution of Siglec-1 in iDCs and mDCs at different incubation times with VLPs. After 5 min of VLP capture, only a very modest colocalization of VLPs with Siglec-1 was observed in both iDCs and mDCs (being slightly higher in the case of mDCs, *figure supplement 5A*). Colocalization progressively increased in time for both types of DCs, but became significantly higher for mDCs at 60 min of VLP incubation (*figure supplement 5A and Supplementary figure S5-source data Fig. S5*). This indicates that Siglec-1 is involved in the initial capture of VLPs by DCs regardless of their activation state. Notably, in contrast to iDCs which showed no significant changes in Siglec-1 clustering during the first 30 min of VLP capture (*Figure 5A and 5B and Figure 5-source data 5*), mDCs already exhibited increased Siglec-1 clustering at these times (*Figure 5C and D*). The latter was also accompanied by an increase in the sizes of Siglec-1 spots colocalizing with VLPs (*figure supplement 5B*). Moreover, cumulative intensities plots of individual VLPs as a function of distance from their center of mass, showed no changes on iDCs for different capture times (*Figure 5E*), whereas a significant increase in the proximity, i.e., polarization, of VLPs in mDC was observed at 30 min (*Figure 5F*). These results reveal remarkable changes in the nano- and meso-scale organization of Siglec-1 upon VLP capture on mDCs occurring already at early capture times. In contrast, more modest changes are observed on iDCs and are only detected after 60 min of VLP capture.

**Figure 5.**
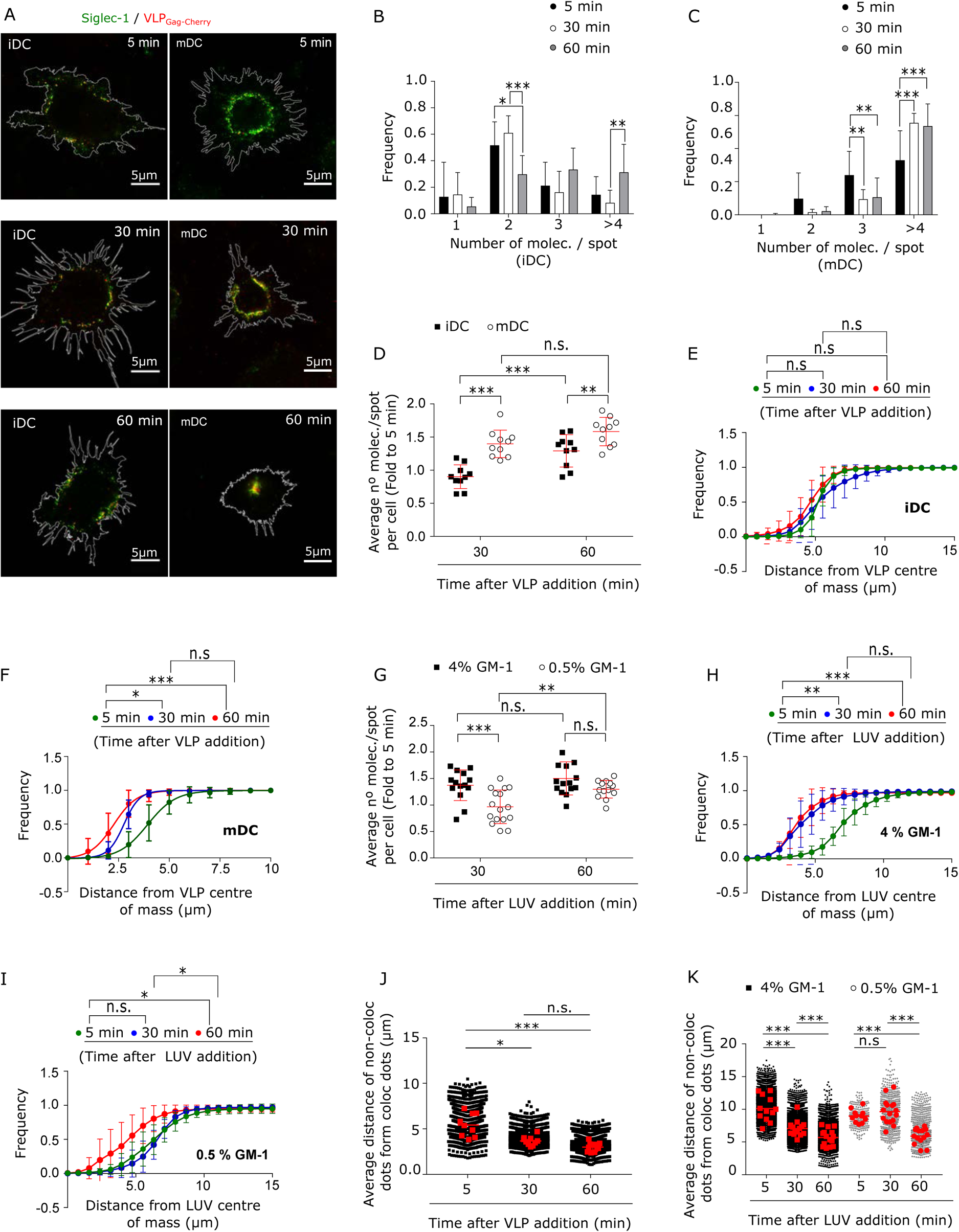
Interaction with HIV-1 particles induces Siglec-1 changes at the nano- and meso-scale towards the formation of the VCC. (A) Representative STED images of iDC (left) and mDC (right) immunostained with anti-Siglec-1 after different times of VLP-Gag-Cherry capture. (B-C) Frequency histograms of Siglec-1 molecules/spot colocalizing with VLP-Gag-Cherry in iDC (B) and mDC (C) at different times of VLP capture. Mean ± S.D. of a minimum of 10 cells per incubation time in iDC and mDC. (D) Fold increase in the number of Siglec-1 molecules/spot with respect to the average value at 5 min. Mean ± S.D. of a minimum of 10 cells per time condition in iDC and mDC. (E-F) Cumulative frequency plots of VLP particles colocalizing with Siglec-1 as a function of the distance from the VLP intensity center-of-mass, in iDC (E) and mDC (F) at different times of VLP capture. Dots show the mean ± S.D. of a minimum of 9 cells per condition and lines show the sigmoidal fit to the data. (G) Same as in D, but in mDC incubated for different times with LUVs at 0.5% and 4% GM-1 concentrations. Mean ± S.D. of a minimum of 13 cells per condition. (H-I) Same as in (E-F) but in mDCs incubated for different times with 4% (H) and 0.5% (I) of GM-1. Dots show the mean ± S.D. of a minimum of 7 cells per time and condition. (J) Box plots showing the distance of non-colocalizing spots to colocalizing Siglec-1/VLP spots in mDC for different incubation times. Small dots show the average distance of the whole population of non-colocalizing spots from each Siglec-1/VLP colocalizing spot, and red squares denote the average of all colocalizing spots in an individual cell (at least 9 cells per time). (K) Same as in (J) but in mDC incubated with 4% and 0.5% GM-1 LUVs (minimum of 11 cells per time and condition). Statistical analysis in the legends of panels (E,F,H,I) corresponds to a one-way ANOVA comparing the distance from the center-of-mass of VLPs (E,F) or LUVs (H,I) at which we recover 50% of the total intensity of VLP-Gag-Cherry or LUVs-tRed.

The fact that VLP polarization occurs concomitantly to the progressive growth of Siglec-1 clusters colocalizing with VLPs on mDCs suggests the possibility that active Siglec-1 clustering, promoted by their interaction with multiple gangliosides carrying sialic ligands in the HIV-1 membrane, regulates the trafficking of the receptor/virus complexes to the VCC. To investigate this scenario, we pulsed mDCs with LUVs carrying low and high GM-1 concentrations, and assessed the evolution of Siglec-1 nanoclusters and LUV spatial dispersion within the plasma membrane of mDCs along time (representative images shown in *figure supplement 5C*). In the case of high GM-1 concentrations (4%), with more ligand available to interact with Siglec-1 receptors, 30 min of binding led on average to ~1.5-fold increase in the number of colocalizing Siglec-1 molecules/spot with respect to the values measured immediately after binding (5 min) (*Figure 5G*). In contrast, in the case of LUVs with low (0.5%) GM-1, the number of colocalizing Siglec-1 molecules/spot did not increase to the same level as to the high GM-1 concentration until 60 min (*Figure 5G*). Interestingly, LUVs with high GM-1 concentration exhibited a clear polarization towards the cell center at 30 min of capture (*Figure 5H*), whereas the low GM-1 LUVs did not show such polarization until 60 min (when the number Siglec-1 molecules/ spot colocalizing with LUVs also starts to increase) (*Figure 5I*). Altogether these results suggest that active Siglec-1 clustering, which is mediated by the sole interaction of Siglec-1 with its ligand GM-1, regulates the trafficking of Siglec-1 receptors towards the formation of the VCC.

Notably, Siglec-1 receptors not colocalizing with either VLPs or high GM-1 concentration LUVs also showed some degree of polarization towards the cell center after 30 min of incubation, which was reflected by a significant increase in the proximity between non-colocalizing and colocalizing spots along time (*Figure 5J and K*). In contrast, such effect was only observed at later times (60 min) when cells were pulsed with low GM-1 concentration LUVs (*Figure 5K*), indicating that at these low concentrations, polarization progressed slower. Finally, we also observed a slight but significant increase in the clustering of non-colocalizing Siglec-1 spots for the cases of VLPs and of high GM-1 concentration LUVs at early capture times, which was only detected at 60 min for low GM-1 LUVs (*figure supplement 5D*). Altogether these results suggest that Siglec-1 enhanced nanoclustering induced by multiple ligand/receptor interactions causes a global nano- and meso-scale rearrangement of the receptor towards its trafficking to the VCC.

### Binding of Siglec-1 to HIV-1 particles induces a global reorganization of the actin cytoskeleton allowing the formation of a single VCC in mDCs

We have shown that Siglec-1 lateral mobility and nanoscale organization at steady-state is controlled by the underlying actin cytoskeleton, dependent on formins and Rho activity. To address how the actin cytoskeleton might also regulate the changes in the nano- and meso-scale organization of Siglec-1 upon binding to gangliosides carrying sialic acid ligands, we fixed and stained mDCs for actin and Siglec-1 after different times of incubation with either VLPs or LUVs carrying high and low GM-1 concentrations (*Figure 6A and Figure 6-source data 6*). After 30 min of incubation, colocalization of Siglec-1 and VLPs or high GM-1 (4%) LUVs occurred in a ring-shaped compartment within the first 4 µm from the basal membrane (*Figure 6A* and *figure supplement 6A and B and Supplementary figure S6-source data Fig. S6*). Moreover, this was accompanied by a clear polarization of the actin cytoskeleton, characterized by a significant reduction of the cell area at the basal planes where the ring was formed (*Figure 6B* for VLP, and *Figure 6C-E* for LUVs) and by the onset of numerous membrane ruffles in the apical parts of the cell (bottom panels in *Figure 6A*). By contrast, pulsing the cells with low GM-1 (0.5%) concentration LUVs, resulted in a significant delay in the accumulation of Siglec-1 and LUVs in a narrow ring-shaped compartment (*Figure 6A*), as well as in the constriction of the basal membrane (*Figure 6C-E*). Together, these results show a direct correlation between the ability of VLPs or LUVs (with high GM-1 content) to promote enhanced Siglec-1 nanoclustering and the induction of downstream global actin rearrangements.

**Figure 6.**
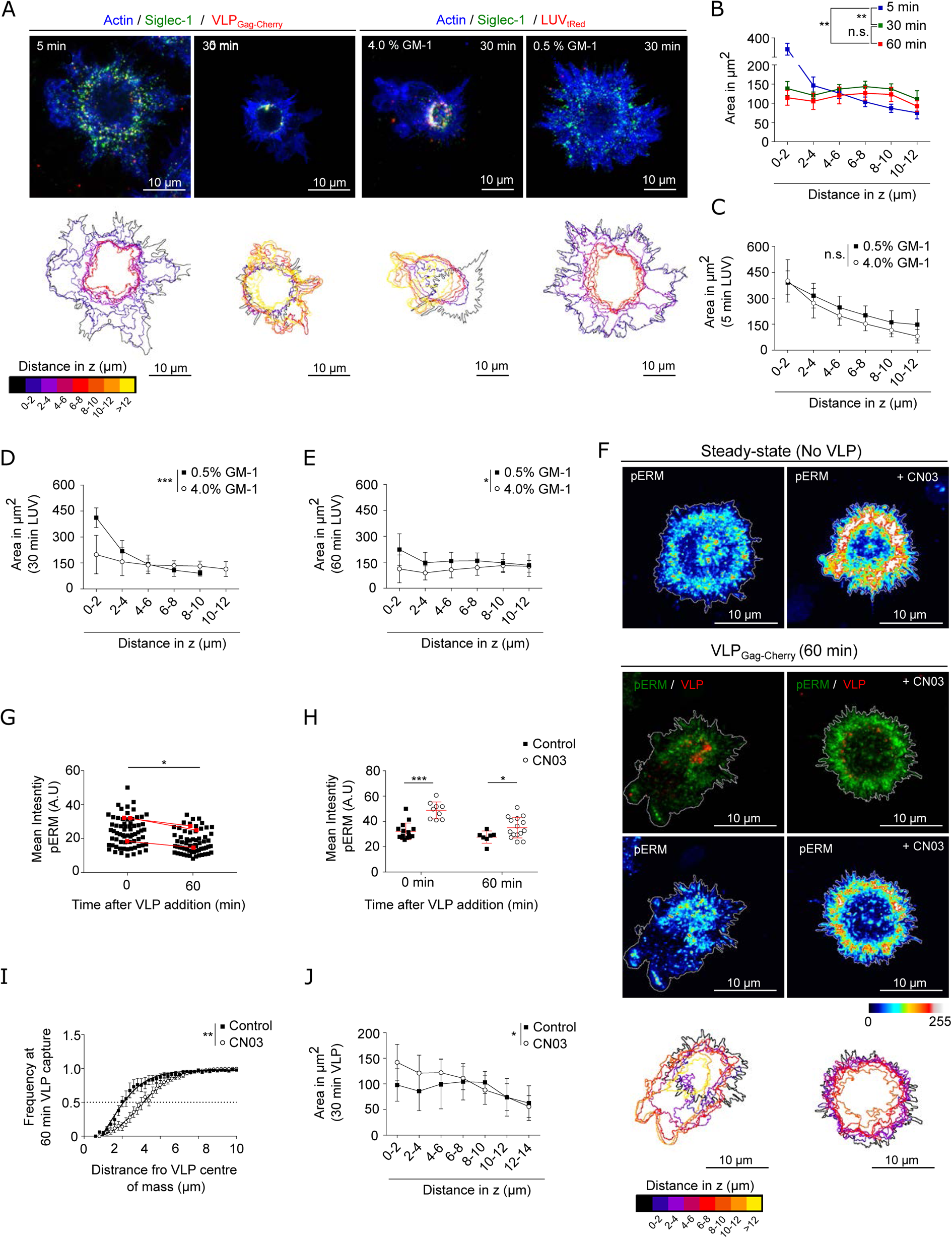
Binding of Siglec-1 to HIV-1 induces a global reorganization of the actin cytoskeleton which allows the formation of a single VCC in mDCs. (A) Confocal maximum intensity projections of fixed mDCs immunostained with anti-Siglec-1, SiR-actin and incubated for different times with VLP-Gag-Cherry or tRedLUVs. Color-coded outlines below each image show the cell perimeter in the axial plane. (B) Cell area in the axial plane in mDC for different incubation times with VLPs. Data correspond to the mean ± S.E.M of 4 (time-points 5 min and 30 min) and 3 donors (60 min), 7 cells/donor and condition. (C-E) Plots as in B for mDC pulsed with LUVs carrying 0.5% and 4% GM-1 for different incubation times (C, 5 min; D, 30 min; E, 60 min). Mean ± S.D. of minimum of 7 cells per time and condition. (F) Confocal projections of mDC stained with anti-pERM in control conditions (left) or pre-treated for 4h with CN03 (right), in the absence (top panels) or the presence of VLP-Gag-Cherry for 60 min (bottom panels). Below scale-colored projections of the cell edges from the bottom to the top of control and CN03 treated mDC after 60 min of VLP capture. (G) Average pERM intensity in mDC before and after 60 min of VLP capture (3 donors, at least 7 cells/donor). Red dots are the mean values of each donor. (H) Average pERM intensity in control mDC and cells pre-treated with the Rho activator CN03 before and after 60 min of VLP capture. Mean ± S.D. of a minimum of 7 cells per condition. (I) Cumulative frequency plots of VLP intensity as a function of the distance from their center of mass in control mDC and cells pre-treated with CN03 after 60 min of VLP capture. Mean + S.D. of a minimum of 7 cells per condition. (J) Cell area in the axial plane in control and CN03 treated mDC after 60 min of incubation with VLPs. Mean ± S.D. of a minimum of 7 cells per condition. Statistics in the legends of panels B, C, D, E, J correspond to the significance of a two-way ANOVA test depending on time after VLP addition (A) percentage of GM-1 (C, D, E) or treatment (control mDC vs CN03 treated cells). Statistics in the legends of panel I correspond to the significance of a Mann-Whitney comparing the distance from the center-of-mass of VLP at which we recover 50% of the total intensity of VLP-Gag-Cherry.

We described that Siglec-1 steady-state distribution was mainly found in regions of the plasma membrane regulated by Rho activity and enriched in pERM. Interestingly, when mDCs were pulsed with VLPs we observed a general reduction of pERM as compared to control (i.e., in the absence of VLP) (*Figure 6F and G*). This drop in pERM suggests a drop in Rho activity and indeed, we observed a similar decay in the levels of the ROCK downstream effectors pMLC (*figure supplement 6C and D*) and p-cofilin (*figure supplement 6E*) after 60 min of VLP capture. Thus, Rho inactivation might be responsible for the major actin reorganization observed and the polarization of VLPs towards the formation of the VCC.

To test this hypothesis, we pre-treated mDCs with the Rho activator CN03. Activation of Rho led to a homogeneous increase in the levels of pERM as compared to control cells, and such difference remained after 60 min of incubation with VLPs (*Figure 6F, H*). Importantly, pre-treatment with CN03 delayed the polarization of VLPs into a ring-shape compartment (*Figure 6I*) and delayed the constriction of the membrane within the planes of VLP accumulation (*Figure 6J*). Altogether these results strongly indicate that actin remodeling associated to Siglec-1 clustering caused by multiple ligand interactions and the subsequent effect on Rho inactivation are essential for the formation of the VCC in mDCs.

## Discussion

Several studies have reported the importance of the actin cytoskeleton in almost all the steps of HIV-1 infection (Audoly et al., 2005; Felts et al., 2010; Gladnikoff et al., 2009; Harmon et al., 2010; Iyengar et al., 1998; Izquierdo-Useros et al., 2011; Jolly et al., 2007; Kerviel et al., 2013; Li et al., 2017; Ménager and Littman, 2016; Nikolic et al., 2011; Sasaki et al., 1995; Shrivastava et al., 2015; Wen et al., 2014). In this study we demonstrated that RhoA activity and formin-dependent actin polymerization regulate Siglec-1 organization on the membrane of mDCs by forming receptor nanoclusters that increase the avidity for HIV-1 capture (*Figure 7A*). Our results indicate that differences in actin polymerization mechanisms between iDCs and mDCs are responsible for clustering of Siglec-1 upon DC activation. Along these lines, recent studies have shown that Arp2/3-dependent branched actin is predominant in iDCs, whereas formation of actin bundles by formins is the main mechanism in mDCs (Vargas et al., 2016). Accordingly, our results show that in mDCs, Siglec-1 exhibits an uneven distribution, with high density regions in polarized areas of the plasma membrane that are enriched in the Rho downstream effector pERM, and can be disrupted by the inhibition of RhoA, ROCK and formins. Interestingly, the abrogation of basal nanoclustering decreases the binding of HIV-1 particles to Siglec-1. Moreover, liposomes with low concentrations of the Siglec-1 ligand GM1 can only bind to large Siglec-1 clusters. These results indicate that the polarized distribution and nanoclustering of Siglec-1 in high density regions of the membrane may provide very efficient docking platforms capable to interact with HIV-1 and other enveloped viruses even with limited concentrations of gangliosides on their membrane (*Figure 7A*). It is tempting to speculate that mDCs subjected to chemokine gradients in 3D environments, which enhance the polarization of Rho and formin actin polymerization at the trailing edge of migrating cells (Lämmermann et al., 2009; Lämmermann and Sixt, 2009; Nitschké et al., 2012; Vargas et al., 2017), could increase HIV-1 capture by enhancing the confinement of Siglec-1 receptors at the back of the cell (the uropod). During the natural course of HIV-1 infection there is an increase in the levels of LPS in the serum, mainly due to bacterial translocation induced by damage in the gut epithelium (Brenchley et al., 2006). It would be interesting to address if the expected changes in the nanoscale organization of activated DCs during migration from peripheral tissues are maintained once DCs reach the lymph nodes, increasing the efficiency of viral capture and *trans-*infection.

**Figure 7.**
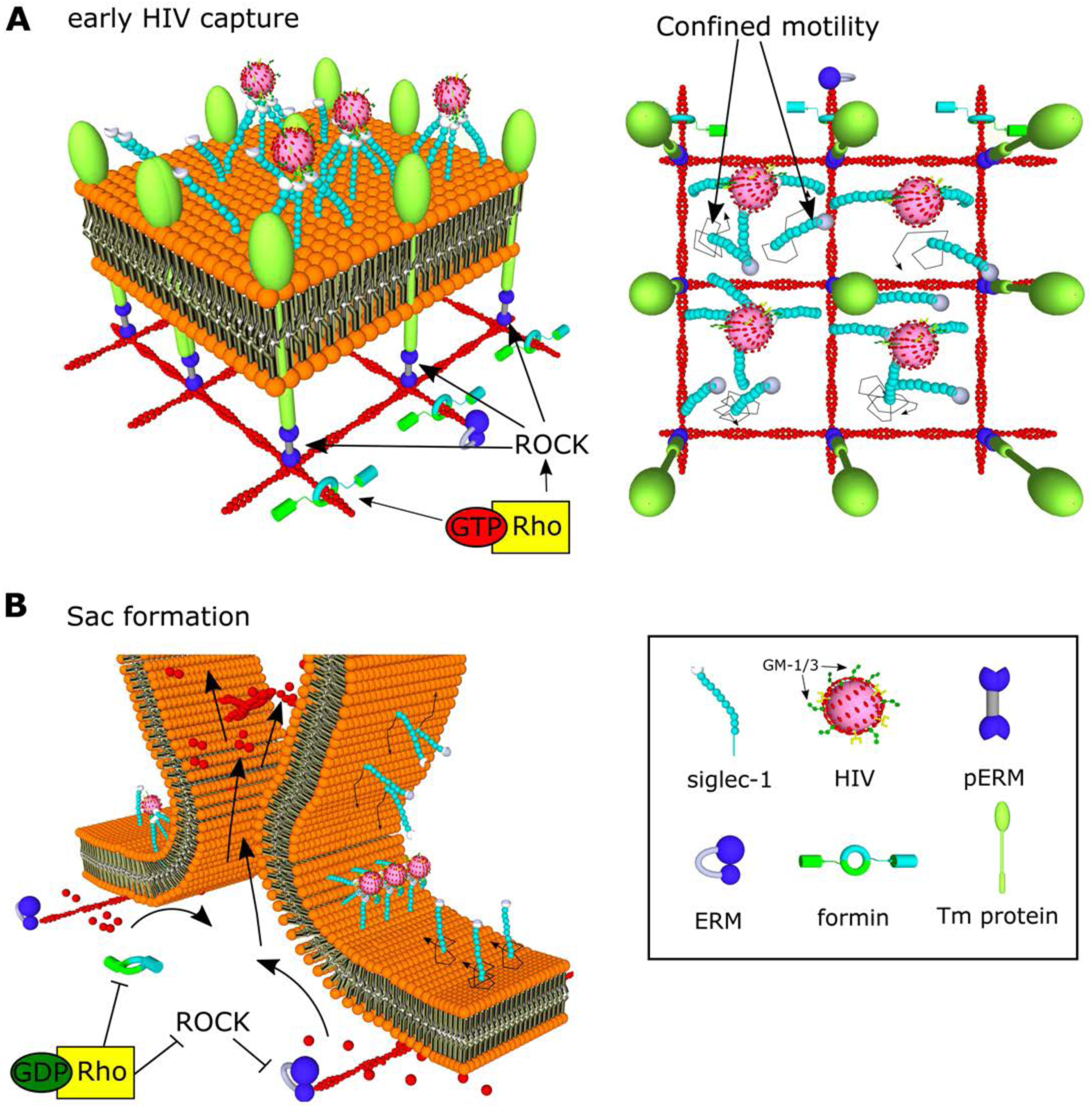
Model of HIV-1 capture by Siglec-1 and polarization towards the formation of the VCC in mDC. (A) In steady state, Siglec-1 receptors in mDCs are organized in small nanoclusters proximal to each other. This basal nanoclustering occurs by spatiotemporal confinement of Siglec-1 diffusion into plasma membrane regions dependent on Rho activity, its downstream effector ROCK, pERM and formin mediated-actin polymerization (see in the middle panel a representation of the cross-section in the 3D reconstruction of a whole cell at early stages of HIV-1 capture). We propose that Siglec-1 mobility is restricted by transmembrane proteins that interact with the filamentous cortical actin cytoskeleton through activated ERM proteins in polarized regions of the mDC membrane (right). Siglec-1 basal nanoclustering is especially relevant to increase the avidity of Siglec-1 for the gangliosides in the membrane of the HIV-1, and therefore regulates the capture capacity of HIV-1 by mDC. (B) At late stages of HIV-1 capture, Siglec-1 clustering increases and most receptors accumulate in a polarized ring-shaped compartment that progressively constrains the membrane (see in the middle panel a representation of the cross-section in the 3D reconstruction of a whole cell at late stages of HIV-1 capture). These changes are accompanied by massive actin rearrangements that include extensive membrane ruffling above the plane in which Siglec-1 and HIV-1 accumulate and by a contraction of the membrane below the ring. This polarization requires a temporal drop in Rho activity. We propose that Rho inactivation would allow for the free diffusion of Siglec-1 through the membrane until being trapped by direct interaction with HIV-1 in the highly dense Siglec-1 regions in which initially most viruses are bound. In parallel, a drop in Rho activity would also lead to actin depolymerization and a decrease in membrane tension together with a retraction of the basal membrane. Both factors would favor the proximity between Siglec-1 bound to HIV-1 clusters and the progressive bending of the membrane until the formation of a single virus sac compartment.

Previous reports have shown that the organization in membrane microdomains of receptors involved in viral binding, like DCSIGN, enhances the avidity of those receptors for their ligands (Cambi et al., 2004). However, this effect is largely dependent on the size of the cargo. Thus, large beads of 1 μm coated with mannose residues, but not small viral particles (~100 nm), can simultaneously engage multiple receptors even in the surface of mildly activated DCs, which contrary to iDCs, do not display a nanoscale organization of DC-SIGN in highly dense microdomains. Siglec-1 is involved in the capture of other cargoes carrying sialylated gangliosides aside from HIV-1 (Perez-Zsolt et al., 2019; Saunderson et al., 2014). Some of them have sizes considerably larger than HIV-1, like the Ebola virus (>1 µm length), whereas others, like exosomes, exosome-like vesicles, ectosomes, or apoptotic vesicles can have a broad range of sizes below 100 nm (Théry et al., 2009). Thus, Siglec-1 nanoclustering may have a different impact in the capture of particulate antigens depending on their structural properties. This possibility can extend the relevance of Siglec-1 spatial organization to other immunological processes like antigen presentation of MHC-I and MHC-II/antigen complexes captured from secreted exosomes (André et al., 2004; Benet et al., 2021; Chaput et al., 2004; Segura et al., 2007; Théry et al., 2002; Wakim and Bevan, 2011).

Our study also underscores the mechanism by which formin-dependent actin polymerization and Rho-ROCK signaling regulates the nanoscale organization of Siglec-1. Our observations on the diffusion behavior of Siglec-1 in mDCs, with a high fraction of molecules displaying immobile or confined motions, suggest that Siglec-1 nanoclustering results from its spatiotemporal confinement in the membrane, which could facilitate the lateral interaction with proximal Siglec-1 receptors. However, Siglec-1 receptors do not possess known cytosolic motifs that directly or indirectly interact with the actin cytoskeleton (Bornhöfft et al., 2018). Nevertheless, it is known that the lateral diffusion of lipids and receptors in the plasma membrane can be constrained by interactions with immobile transmembrane proteins or “pickets”, directly connected to the cortical actin through linker proteins like ezrin (Freeman et al., 2018; Kalay et al., 2014; Kusumi et al., 2012; Trimble and Grinstein, 2015). The fact that pharmacological inhibition of ROCK and its downstream effector ERM can partially disrupt Siglec-1 nanoclustering and its polarization in highly dense areas of the plasma membrane supports the existence of such protein pickets acting on Siglec-1 (*Figure 7A*). Proteins like CD44 can exert this picket function and confine the diffusion of other transmembrane receptors at zones enriched in pERM and formin-dependent actin polymerization at the trailing edge of mDCs (Freeman et al., 2019; Sil et al., 2020). It remains to be investigated whether amongst the cofactors needed for the formation of the VCC [19], there are also proteins acting as diffusion barriers to confine Siglec-1 in restricted areas of the cell.

Lastly, we showed that active Siglec-1 clustering promoted by interactions with HIV-1 is essential to regulate its trafficking towards the formation of the VCC. Importantly, the proximity between VLPs proceeds in parallel with a progressive clustering of Siglec-1, which is much more pronounced in the case of mDCs as compared to iDCs, and dynamically dependent on the concentration of gangliosides in the viral membrane. Indeed, when mDCs are pulsed with liposomes carrying low concentrations of GM-1 we observed a significant delay in the growth of Siglec-1 clusters and a more disperse distribution of these liposomes within the cell membrane, as compared to those with high GM-1 concentrations. In addition, the polarization of viral particles in a ring-shape compartment triggered several morphological changes in mDCs: emergence of membrane ruffles, shrinkage of the basal membrane, and constriction of the cell membrane at the plane where VLPs accumulate. These results indicate that active clustering of Siglec-1 is capable of triggering a downstream signaling cascade that dramatically affects the actin cytoskeleton of mDCs. Accordingly, we showed that VLPs accumulation into the ring-shape compartment coincides with a decay in the levels of the ROCK effectors pERM, pMLC and p-cofilin. Moreover, pharmacological activation of Rho caused a delay in the constriction of the ring-shape compartment, indicating that Rho inhibition is required for the proper polarization of VLPs towards the formation of the VCC. A decay in Rho activity may, on one hand, decrease actin polymerization by formins and activate severing of filamentous actin by cofilin (Sit and Manser, 2011), which could explain the retraction of the basal membrane. On the other hand, the inhibition of pERM, one of the main regulators of membrane tension (Y. Liu et al., 2012; Rouven Brückner et al., 2015) might facilitate the observed constriction of the membrane at the plane where the virus accumulate. Both phenomena would enhance the proximity between Siglec-1/HIV-1 complexes facilitating the lateral fusion of molecules until a VCC is formed (*Figure 7B*).

How Siglec-1 clustering is capable to activate such changes needs further investigation. The fact that the cytosolic domain of Siglec-1 is not essential for the formation of the VCC reinforces the idea that other cofactors might co-cluster with Siglec-1 to activate intracellular signaling cascades. Interestingly, the cross-linking of several tetraspanins, which are highly enriched in the VCC [13,14,16], can induce actin rearrangements through the modulation of Rho GTPases (Brazzoli et al., 2008; Delaguillaumie et al., 2002; Jones et al., 2016; W. M. Liu et al., 2012). Furthermore, the spatial organization of tetraspanins in the so-called tetraspanin-enriched microdomains (TEM) is also regulated by interactions with certain lipids like cholesterol (Zimmerman et al., 2016) and gangliosides (Odintsova et al., 2006). Yet, it is not known if some trans-interactions between the tetraspanin web and the gangliosides exposed in the membrane of HIV-1 can also affect the coalescence of multiple TEM (Delaguillaumie et al., 2004). The formation of the VCC could be the result of a synergistic cooperation between the capacity of several tetraspanins to modulate positive and negative membrane curvature (79,80) and the differential lipid composition of TEMs, which could facilitate domain-induced inward budding to minimize the line energy at the TEM-membrane interface (Baumgart et al., 2003; Lipowsky, 1992; Liu et al., 2006). In this regard, the use of single particle tracking and super-resolution microscopy can be of great interest to further investigate the molecular composition of Siglec-1/HIV-1 clusters within the course of the VCC formation, as well as to identify new key regulators of this process.

## Materials and Methods

### Antibodies and reagents

Mouse monoclonal antibodies to Siglec-1 (Hsn 7D2) and rabbit polyclonal antibodies against cofilin and phosphor-cofilin (phospho S3) were obtained from Abcam. Rabbit antibodies against pERM (48G2) and pMLC were purchased from Cell Signaling. Anti-Mouse IgG F(ab) ATTO488 was from Hypermol and Anti-Rabbit IgGF(ab) Cy3 was from Jacson Immunoresearch. The SIR-Actin Spirochrome Kit (CY-SC001), the Rho Inhibitor I ADP ribosylation of Rho Asn-41 (CT04) and Rho Activator II (CN03) were obtained from Cytoskeleton. The phosphor-ezrin inhibitor NSC668394 was from Calbiochem. The pan-formin inhibitor SMIFH2, the Arp2/3 inhibitor CK-666 and the actin polymerization inhibitor CytoD were purchased from SIGMA. The ROCK inhibitor Y-27632 was obtained from Merk Millipore.

### Primary cell cultures

Peripheral blood mononuclear cells (PBMC) were obtained with a Ficoll-Hypaque gradient (Alere Technologies AS) from HIV-1-seronegative donors, and monocyte populations (>97% CD14+) were isolated with CD14-positive selection magnetic beads (Miltenyi Biotec). DCs were obtained by culturing these cells in the presence of 1,000 IU/ml granulocyte-macrophage colony-stimulating factor and interleukin-4 (both from R&D) for seven days and replacing media and cytokines every two days. Activated DCs were differentiated by culturing iDCs at day five for two more days in the presence of 100 ng/ml lipopolysaccharide (LPS, Sigma-Aldrich) to induce Siglec-1 expression.

### Plasmids and viral stocks

HIV-1-Gag-mCherry-env HXB2 were obtained by co-transfection 1:1 (w/w) of the molecular clones pcDNA HIV Gag Cherry (kindly provided by Dr. Gummuluru; 15 µg) and pcDNA HIV-1 env HXB2 (15µg). HEK-293T cells were transfected with calcium phosphate (Clontech) in T75 flasks using a total of 30 μg plasmid DNA. 72 hours post-transfection, supernatants containing virus were cleared of cellular debris by centrifugation, filtered (0.45 μm; Millipore) and frozen at −80 °C until use.

### Generation of LUVs

LUVs were prepared following the extrusion method described in (Mayer et al., 1986). Lipids were purchased from Avanti Polar Lipids and gangliosides were obtained from Carbosynth. The LUV-tRed lipid composition was: POPC 25 mol%: 1,2-dipalmitoyl-sn-glycero-3-phosphocholine (DPPC) 16 mol%: brain sphingomyelin (SM) 14 mol%: cholesterol (Chol) 39 mol % : GM-1 4 to 0.25 mol%. All the LUVs contained 2 mol% of 1,2-dihexadecanoyl-sn-glycero-3-phosphoethanolamine (DHPE)-tRed (Molecular Probes). Lipids were mixed in chloroform: methanol (2:1) and dried under nitrogen. Traces of organic solvent were removed by vacuum pumping for 1–2 hours. Subsequently, the dried lipid film was dispersed in HEPES-sodium buffer and subjected to ten freeze-thaw cycles prior to extruding ten times through two stacked polycarbonate membranes with a 100-nm pore size (Nucleopore, Inc.) using the Thermo-barrelextruder (Lipex extruder, Northern Lipids, Inc.). To perform mDC pulse with equal concentrations of LUV displaying similar fluorescence intensities, tRed containing LUVs concentration was quantified following the phosphate determination method (Böttcher et al., 1961) and the fluorescence emission spectra was recorded setting the excitation at 580 nm in a SLM Aminco series 2 spectrofluorimeter (Spectronic Instruments).

### Immunoblot

DCs derived from PBMCs treated with LPS for 48 hours were seeded at a concentration of 0.5×10^6^ cells well in 12-well plates coated with poly-L-lysine (30 µg/mL). After 1 hour of adhesion cells were pulsed with 50 µL of VLP-Gag-Cherry particles in each well, and gently scrapped on ice after different times of incubation at 37 °C. Cell pellets were lysed in Triton X-100 lysis buffer (ThermoFisher) supplemented with protease inhibitors (Roche), and 10µg of protein samples were prepared adding NuPAGE LDS Sample Buffer (ThermoFisher) with 2.5% beta-mercaptoethanol (SIGMA). Samples were further denatured at 80 °C for 5 minutes and loaded in 12% acrylamide gels (Bio-Rad). Electrophoresis was carried in Tris-Acetate SDS running buffer (Bio-Rad) at 100V and transfer to nitrocellulose membranes was done in Towbin buffer (25 mM Tris, 192 mM glycine, 20% (v/v) methanol) for 90 minutes at 100V. Membranes were incubated with primary and HRP-secondary antibodies in TBST buffer with 5% milk (RT for 1 hour or overnight at 4 °C) and revealed with an ECL solution (BIO-Rad) following manufacturer’s instructions.

### Generation of single chain antibodies

Monovalent anti-human Siglec-1 Abs were prepared by reduction with DTT at 10mM for 30 minutes at RT. DTT was removed dialyzing the samples in PBS using Zeba™ Spin Desalting Columns (Thermo Scientific) and when required monovalent antibodies were further concentrated using Centricon concentrators according to manufactureŕs instructions. To verify the generation of single chain antibodies we prepared samples of whole and reduced antibodies with NuPAGE LDS Sample Buffer (4x) and run them in NuPAGE™ 7% Tris-Acetate Protein Gels (Bio-Rad) in denaturing, non-reducing conditions. Electrophoresis was performed in Tris-Acetate SDS running buffer (Bio-Rad) at 100V and gels were stained with Coomassie for 1 hour at RT.

### Sample preparation for STED and confocal imaging in fixed cells

1.5×10^5^ immature and LPS (100 ng/mL for 48h) matured DCs were plated on glass slides coated with poly-lysine (10 µg/mL). After DCs adhesion to culture plates, cells were pulsed with either VLP-Gag-Cherry particles (30 µl per 1.5 ×10^5^ cells) or LUVs carrying different concentration of GM-1 (200 µM) for 5 minutes to 1 hour at 37°C. After extensive washing, cells were fixed in PFA 4% (10 min RT) and rinsed with NH_4_Cl (50 mM in PBS). When needed, permeabilization was done with saponin 0.01% in PBS (10 min, RT). Immunostaining was done incubating primary and secondary antibodies (1 hour, RT) in PBS 1% BSA. SIR-actin was added at 1 µM together with primary antibodies. After several washes in PBS glass slides were covered with Fluoromount aquose mounting media (Sigma). When indicated cells were incubated with CytoD (2 µg/mL), CK666 (100 µM), SMIFH2 (25 µM) and Y-27623 (30 µM) for 1 hour at 37 °C in complete RPMI (10% FBS, L-glutamine) prior fixation. NSC668394 (250 µM) was added for 3 hours at 37 °C in complete RPMI. CT04 (2 µg/mL) and CN03 (2 µg/mL) were added for 3-4 hours at 37 °C in RPMI without FBS. When cells were incubated with VLP-Gag-Cherry the concentration of the drugs was maintained for all the time until fixation.

### STED imaging

STED super-resolution images of DCs were acquired with a confocal microscope (Leica TCS SP8, Leica Microsystems) equipped with an oil immersion objective (HCX PL APO CS ×100, Leica) with a 1.4 numerical aperture. Samples were excited with a white light laser (WLL) at 488nm (Atto488), 587nm (mCherry), 583nm (tRed), 633nm (SiR) fixing the laser power to optimal conditions (5-20%) for each experiment. Emission spectra was in the range of 500-540 nm (Atto488), 550-650nm (mCherry and tRed), 645-661nm (SiR). STED laser beams intensities at 592 nm (Atto488) and 775 nm (mCherry, tRed and SiR) were set to 50% and 100% of their power respectively. Images were acquired with a format of 1024×1024 pixels at 400Hz with a pixel size of ~14 nm, adjusting frame accumulation (1-5) and line average (1-5) to a non-saturating signal for each staining, keeping the same conditions for all the cells analyzed in one experiment.

### Quantification of number of molecules per spot

To define the area of Siglec-1 individual spots STED images were processed using Fiji to apply a subtraction of the background and a Gaussian blur filter (sigma radius 1) followed by a difference of Gaussian (smaller/greater sigma 1:3). Then an intensity threshold was used to create binary masks of the individual spots from which we obtain the mean intensity values per each spot within the original images. Thereafter, we used a MATLAB custom code to fit the distribution of intensities of individual spots from antibodies on glass to a lognormal function (f_1_).

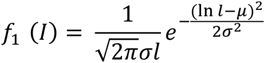

This model corresponds to the expected theoretical distribution for the intensity corresponding to the detection of a fluorescent emitter (Cella Zanacchi et al., 2017; Moertelmaier et al., 2005; Schmidt, 1996). The intensity values obtained from spots on glass were used to define the *µ* (mean) and *σ* (standard deviation) of the lognormal distribution, through its fit to a linear combination of N = 2 functions. These parameters were used as a single molecule reference to define the stoichiometry of the fluorescence of Siglec-1 receptors in the cells measured under identical experimental conditions (Torreno-Pina et al., 2016; Van Zanten et al., 2009). As expected, the intensity histograms of Siglec-1 spots on samples showed higher intensities and broader distributions than the antibodies on glass, indicative of a mixture of different populations of nanoclusters composed by a different number of molecules. To calculate the probability distribution of molecules per spot, the intensity histograms of Siglec-1 spots on cells was fitted to a model distribution g_N_ (I) composed of a linear combination of functions as described in (Martínez-Muñoz et al., 2018).

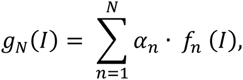

Where *f_n_* shows the intensity distribution intensity of a spot containing n receptors, and *α_n_* is the relative weight of this distribution so that 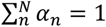, being *N* the maximum number of receptors (in our analysis 12) (Moertelmaier et al., 2005). We considered that the distribution for a spot containing *n* receptors could be obtained recursively as

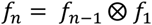

where ⊗ represents the convolution of the intensity distribution lognormal functions for n = 1,2,…,12 (Moertelmaier et al., 2005; Schmidt, 1996; Torreno-Pina et al., 2016).

### Determination of number of molecules/spot colocalizing with VLPs

To classify Siglec-1 clusters according to their colocalization with VLPs, STED images from each channel were processed as described in quantification of molecules per spot followed by an intensity threshold segmentation of individual spots to create binary masks. The binary images of segmented spots in each channel were overlapped to identify Siglec-1 colocalizing clusters (with an average area overlap for each siglec-1 spot of ~ 60% ± 27%). Then, we obtained the Siglec-1 mean intensity values of the spots colocalizing with VLPs from the original unprocessed images, which we used to calculate the probability distribution of number of molecules per spot in the Siglec-1/VLP colocalizing fraction as described above (see quantification of the number of molecules per spot).

To quantify the local density of Siglec-1 molecules surrounding the colocalizing spots, we used a custom MATLAB code to select all the spots whose centroids were enclosed in different radius from the centroid of each colocalizing spot. We then used their mean intensity values to assign a number of molecules (*n*) to each spot (see quantification of molecules per spot). The density (*d*) of molecules within a radius *r* was,

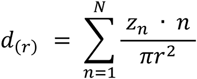

Where *Z_n_* is the total number of spots with *n* molecules and *N* the maximum number of molecules per spot.

To quantify the dispersion of VLPs and LUVs colocalizing with Siglec-1 we measured the distance of each individual particles from its center to the center-of-mass of all the Siglec-1 spots. This distance *vs* the accumulated intensity frequency was fit to a sigmoidal equation from which we obtained the distance covering 50% of the Siglec-1 intensity for each cell, which was used as reference parameter for statistical analysis. To quantify the proximity between Siglec-1 spots colocalizing or not with VLP/LUV, individual Siglec-1 spots were segmented for colocalization as described above and we measured the average distance between the centers of all non-colocalizing spots to the center of each individual colocalizing spot.

### Generation of *in-silico* random images

To generate *in-silico* images with a random distribution we calculated the FWHM of individual spots by averaging the intensity line profiles of multiple spots on glass and fitting the data to a Gaussian function. In this way we generated *in-silico* spots with a PSF and an amplitude equivalent to the values obtained from the Gaussian fits of the real data. For each cell the total number of Siglec-1 molecules randomly distributed in a defined cell area was calculated according to the total number of spots (*Z*) quantified in the experimental data and the probability distributions (*α_n_*) of *n* molecules per spot up to a maximum of *N*=12 molecules per spot.

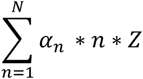

### Analysis of actin and Siglec-1 distribution

To correlate the intensities of Siglec-1 and cortical actin, we processed STED images for Siglec-1 as described in “Quantification of molecules per spot”. Then we measured the mean intensity profiles for each channel from the cell center to the edges in concentric radius of 1.3 pixels. Distance from the cell center were normalized as a percentage of the maximum cell radius for each cell, and intensities at each radius were normalized to the sum of all the intensities for all the radius.

### Under-agarose experiments

Under-agarose chemotaxis assays were performed as described elsewhere (Kopf et al., 2020; Lämmermann et al., 2009). Briefly, we added ultrapure agarose (Invitrogen) at 4% to RPMI supplemented with 20% FBS and Hank buffered solution (GIBCO) at a 1:2:1 ratio. We poured 1 mL of the mixture on 35 mm glass coverslips previously coated with 30 µg/mL of fibronectin. After polymerization at RT we added 4×10^5^ cells in 30 µL of RPMI between the agarose pad and the glass. We allowed the cells to equilibrate for about 15-30 min and then added on the borders of the agarose pad 1 mL of complete RPMI with 2.5 µg/mL of CCL19. After 90 min at 37 °C we directly added to the cells 1 mL of PFA at 8% for 10 min. We gently lifted the agarose pad and processed the sample for immunostaining with SIR-actin and anti-Siglec-1. Two color STED images acquired as previously described were analyzed for the distribution of actin and Siglec-1 staining at the rear and the cell front. Briefly, we defined a 0 angle between the cell center and the center-of-mass of the uropod at the trailing edge, and quantified the percentage of Siglec-1 and actin staining in 40° binning. For the analysis of Siglec-1 molecules per spot, the receptors counted at the rear were all the spots at ±20° (from the 0 angle) and between 140-220° for the front.

### Confocal imaging

Confocal images on fixed DCs were obtained with a confocal microscope (Leica TCS SP8, Leica Microsystems) as specified in “STED images”. Images were acquired with a format of 1024×1024 pixels at 400 Hz with the pinhole set at 1.0 A.U. The whole surface of cells was covered in stacks of 10-15 images from the bottom to the top. We applied intensity thresholds for each staining and calculated the mean intensity values in axial planes of 1.5 µm. For each cell the minimum mean intensity value within the stack was used to normalize the axial distribution of each staining. To measure the area of cells at different axial planes after VLP/LUV capture, we delimited the edges of cells stained with SiR-actin applying a binary mask for each plane and quantified the area enclosed in such perimeter.

### Sample preparation and analysis of Siglec-1 particle tracking

1×10^6^ immature and LPS (100 ng/mL for 48 hours) matured DCs in suspension were incubated in complete RPMI supplemented with 10% FBS and 1 µg/mL of single chain Siglec-1 antibodies on ice for 30 min, after which cells were washed 3 times in PBS and resuspended in 500 µL of complete RPMI. Next, cells were seeded on 35 mm glass coverslips (Corning) coated with poly-L-lysine at 30 µg/mL and after 30-60 min of adhesion at 37° C cells we added Fab-Atto488 anti-mouse antibodies at 1 µg/mL for 5 min. Cells were washed several times in PBS and covered with 1 mL of complete RPMI supplemented with 25 mM of HEPES.

Live imaging was done with a confocal microscope (see STED imaging) connected to a temperature control device at 37°C. For the detection of Fab-Atto488, samples were excited with a WLL laser at 488 nm (20% power) setting the emission range at 509-595 nm. Scan laser speed was set at 1000 Hz to record 3 frames per second in a format of 512×512 pixels.

The particle tracking analysis was done using the MOSAIC tool of Fiji (Sbalzarini and Koumoutsakos, 2005). The implementation of the plugin requires the approximate particle radius *w* (in our case 3 pixels), the intensity percentile *r*, which is used to select among the local maxima pixels in a radius *w* the ones that have an intensity above the selected *r* value (in our case the upper 0,5 percentile), the cut-off intensity score *Ts* to remove those pixel values below a intensity threshold (in our case set to 0), the maximum displacement in pixels between contiguous frames (in our settings 5 pixels), and the number of future frames that can be considered to re-connect particles (in our case 2, which means that a single particle can disappear for just one frame to be re-connected with another particle in the next frame).

Individual trajectories of a minimum of 20 points were analyzed calculating their moments of displacement µ_v_, which for a trajectory of a length M_l_ and a frame shift Δn, corresponding to a time shift δt = ΔnΔt, is defined as:

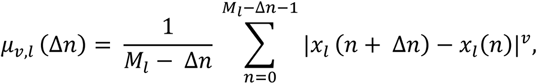

Where x*_l_* is the position vector (x‘(n), y‘(n)) on an individual trajectory *l* at time nΔt. To quantify the particle motion these moments are calculated for ν = 0, 1, 2, …, 6 and Δn = 1,…, M_1/3_,. For all the moments of displacement the scaling coefficients γ_ν_ are determined by a linear least squares regression to lnµ_ʋ_ versus lnδt. The regular diffusion constant for ν = 2 (D_2_) was obtained from the y_0_ intercept of the regression line such as *D*_2_ = 4^−1^ ∗ *e^yo^*. To calculate the fraction of immobile particles we measured the D_2_ values of antibodies on glass acquired in the same imaging conditions as in the cells. We defined mobile particles as those whose D_2_ values were above the 90 percentile of the D_2_ values measured for the antibodies on glass (in our case 0.002 um^2^/s). The slope of the plot of γ_ν_ versus ν, called moment scale spectrum (MSS) was used to classify trajectories according to their motions in confined (<0.2), subdifussive (>0.2-<0.5) and free (>0.5).

## Statistical Analysis

All results were analyzed using GraphPad PRISM 7.0 (ns, p > R 0.05, * p % 0.05, ** p % 0.001; *** p % 0.0001).

Paired T-test analysis were used in figures 1C, 1D, 1G, 2F, 2G, 2K,6G, and S6E.

Non parametric Mann-Whitney tests were used in figures S1F, S1G, S1H, 3H, 3I, S3A, S3B, S3C, S3E, S3J, S3M, 4B, 4C, 4D, S4B, S4C, 6I, S6D.

Kruskal-Wallis with Dunn’s multiple comparison tests were used in figures 4F, 4G, 4J, 5E, 5F, 5H, 5I, 5J.

Two-way ANOVA with multiple comparison Bonferroni tests were used in figures 1B, 1H, 1J, 2H, 2I, 2L, S2C, S2E, S2I, 3B, 3C, 3F, 3J, 3K, S3F, S3G, S3H, S3K, S3L, S3N, 4H, 5B, 5C, 5D, 5G, 5K, S5A, S5B, S5D, 6B, 6C, 6D, 6E, 6H, 6J.

## Acknowledgements

The authors would like to thank K. Borgman and C. Martinez-Guillamon for initial experiments, J. Andilla and M. Marsal for technical support at the Super-resolution Light Nanoscopy (SLN) facility at ICFO. The research leading to these results has received funding from the European Commission H2020 Program (grant agreement ERC Adv788546 (NANO-MEMEC) (to M.F.G.-P.) and Marie Sklodowska-Curie grant 754558-PREBIST (to N.M.)), Government of Spain (Severo Ochoa CEX2019-000910-S, State Research Agency (AEI) (PID2020-113068RB-I00 / 10.13039/501100011033 (to M.F.G.-P.), PID2019-109870RB-I00 (to J.M-P.), (PID2020-117405GB-100 to M.L.), RYC-2017-22227 (to F.C.), RYC-2015-17896 (to C.M.), and PID2019-106232RB-I00/10.13039/501100011033 (to F.C.)), Fundació CELLEX (Barcelona), Fundació Mir-Puig and the Generalitat de Catalunya through the CERCA program and AGAUR (Grant No. 2017SGR1000 to M.F.G.-P.). N.I-U. is supported by grant PID2020-117145RB-I00 from the Spanish Ministry of Science and Innovation. J.M-P an N.I-U are funded by the CIBER de Enfermedades Infecciosas.

## Legends source data

**Figure 1-source data 1:** Excel file containing the source data for Figures 1B-D and Figures 1G, H.

**Figure 2-source data 2:** Excel file containing the source data for Figure 2B and Figures 2D-L.

**Figure 3-source data 3:** Excel file containing the source data for Figures 3B and C, Figures 3F-K

**Figure 4-source data 4:** Excel file containing the source data for Figures 4B-D, Figures 4F-H and Figure 4J.

**Figure 5-source data 5**: Excel file containing the source data for Figures 5B-K

**Figure 6-source data 6:** Excel file containing the source data for Figures 6B-E, and Figures 6G-J

## Supplementary Information

**Figure supplement 1.**
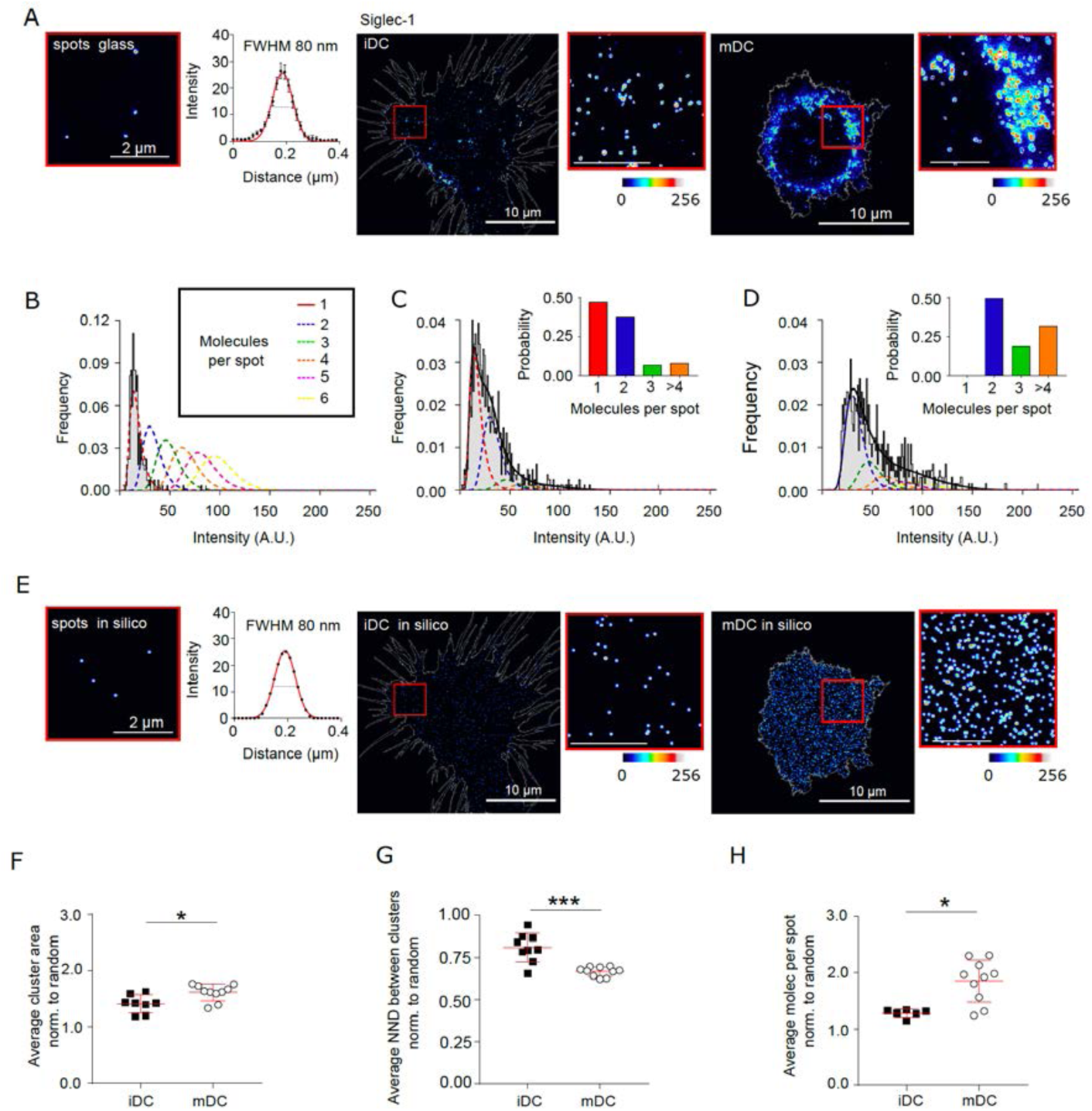
(A) STED images of Siglec-1 antibodies non-specifically adhered to glass and labeled with Fab-Atto488 secondary antibodies with a corresponding Gaussian fit of the intensity (left); and representative STED images of Siglec-1 on iDCs and mDCs together with enlarged views. (B) Histogram of the spot intensities for Siglec-1 antibodies adhered on glass. The black line (and shaded region) shows the lognormal fit of the experimental data, and the dashed lines correspond to the calculated intensity distribution functions for different numbers of molecules per spot. (C-D) Intensity distributions of Siglec-1 spots from the representative images of iDC (C) and mDC (D). Black lines are the fit of experimental intensity distributions from which we extract the probability of having 1 up to > 4 molecules per spot (see bar graphs in the insets). (E) *In-silico* spots generated with a FWHM as extracted from the experimental data shown in panel A for spots on glass (left), Siglec-1 staining in iDC (middle) and mDC (right). To generate the *in-silico* images we quantified the total number of Siglec-1 receptors by taking into account the number of molecules per spot and total number of spots. The resulting number of molecules was randomly distributed within the cell area as spots that were convoluted with the FWHM of experimental data (60-80 nm). (F-H) Average area of Siglec-1 spots per cell (F), average nearest-neighbor-distance (NND) between Siglec-1 spots per cell (G) and average number of molecules per spot per cell (H), in iDC and mDC. For each cell analyzed the values are normalized to the respective *in-silico* values of Siglec-1 random distributions (mean + S.D. of one representative experiment; N=3).

**Figure supplement 2.**
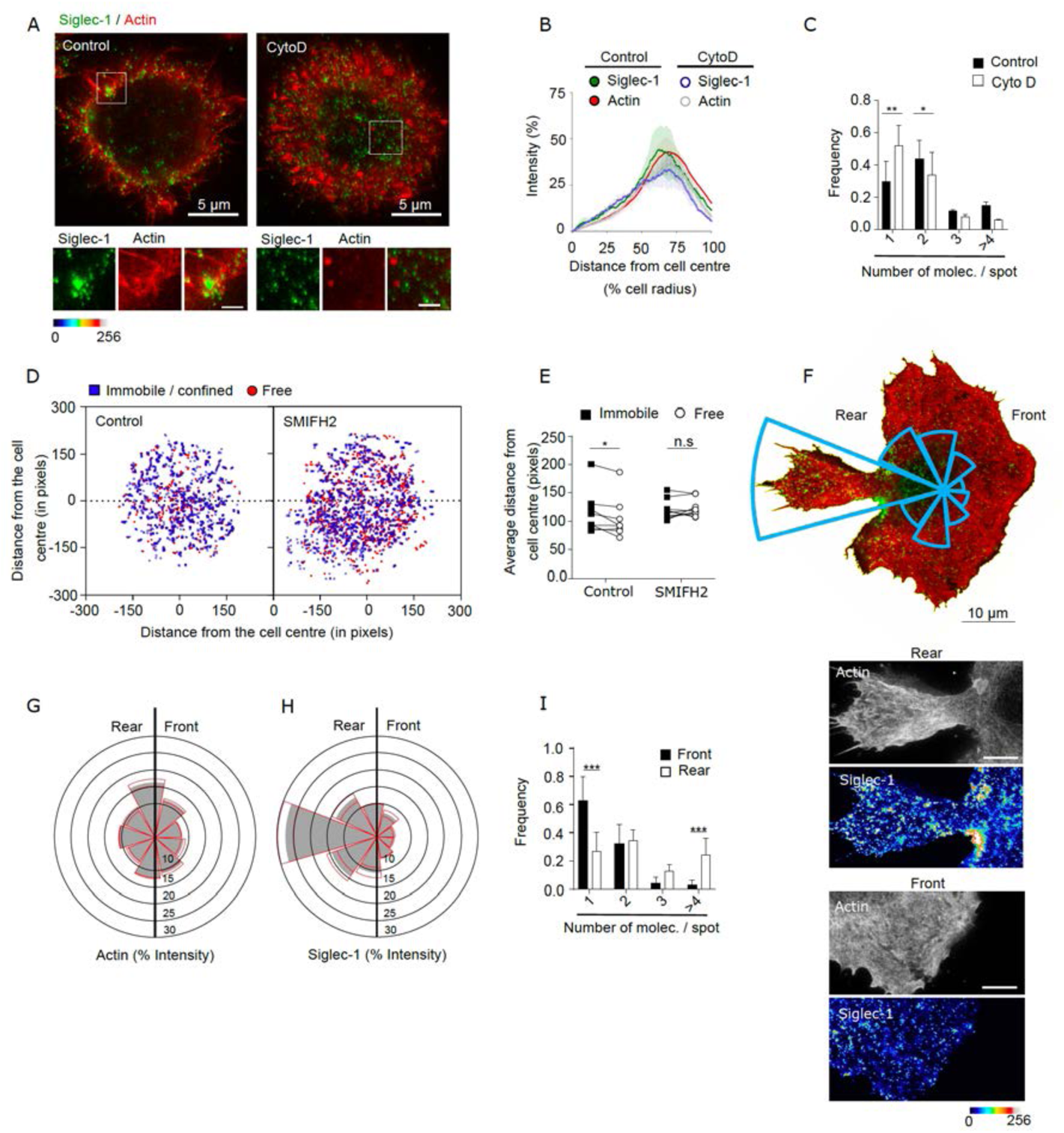
(A) Representative dual color STED images of mDC stained with Siglec-1 and SiR-actin in control conditions and after 1 h treatment with Cytochalasin D (Cyto D, 2 µg/mL). The bottom panels correspond to magnified images of the dashed squares (scale bars: 2 µm). (B) Siglec-1 and actin intensity from the cell center towards the edge expressed as percentage of total intensity for each marker in iDC control mDC and cells treated with Cyto D. Lines represent the mean + S.E.M of two different donors in which a minimum of 10 cells per donor and condition were analyzed. (C) Frequency histogram of the number of Siglec-1 molecules/spot in control mDCs and cells treated with Cyto D. Mean + S.E.M of a minimum of 9 cells per donor from two donors. (D) Plots showing the center mass of individual Siglec-1 trajectories (average *x,y* position in all the frames of a trajectory) in control mDC and cells treated with SMIFH2, colored-coded according to their mobility. The graph shows the centers of all the trajectories analyzed for a minimum of 8 cells per condition from one representative experiment (N=3). (E) Distance from the cell center of immobile/confined and sub-diffusive/free trajectories in control mDC and cells treated with SMIFH2. Black squares show the average distance from the cell center of all immobile/confined trajectories in one cell, and are paired to the average values of all sub-diffusive/free trajectories within the same cell (empty circles). (F) Representative STED image of a mDC fixed and stained for Siglec-1 and SiR-actin after injection between a glass coverslip and a layer of agarose with CCL19 at 2.5 µg/mL. The rose plot histogram shows the distribution of Siglec-1 staining from rear to front in 40° bins. On the bottom magnified insets of Siglec-1 and actin at the rear and the front of the cell (scale bar: 2 µm). (G, H) Rose plot histograms of actin and Siglec-1 intensity distributions on the front and rear end of mDCs subjected to a CCL19 homogeneous gradient in an under-agarose assay. Graphs show the mean + S.E.M (red lines) of two donors with a minimum of 10 cells per donor. (I) Frequency histogram of the number of Siglec-1 molecules/ spot at the rear and the front of mDCs treated as in G. Mean + S.E.M of a minimum of 10 cells per donor from two donors.

**Figure supplement 3.**
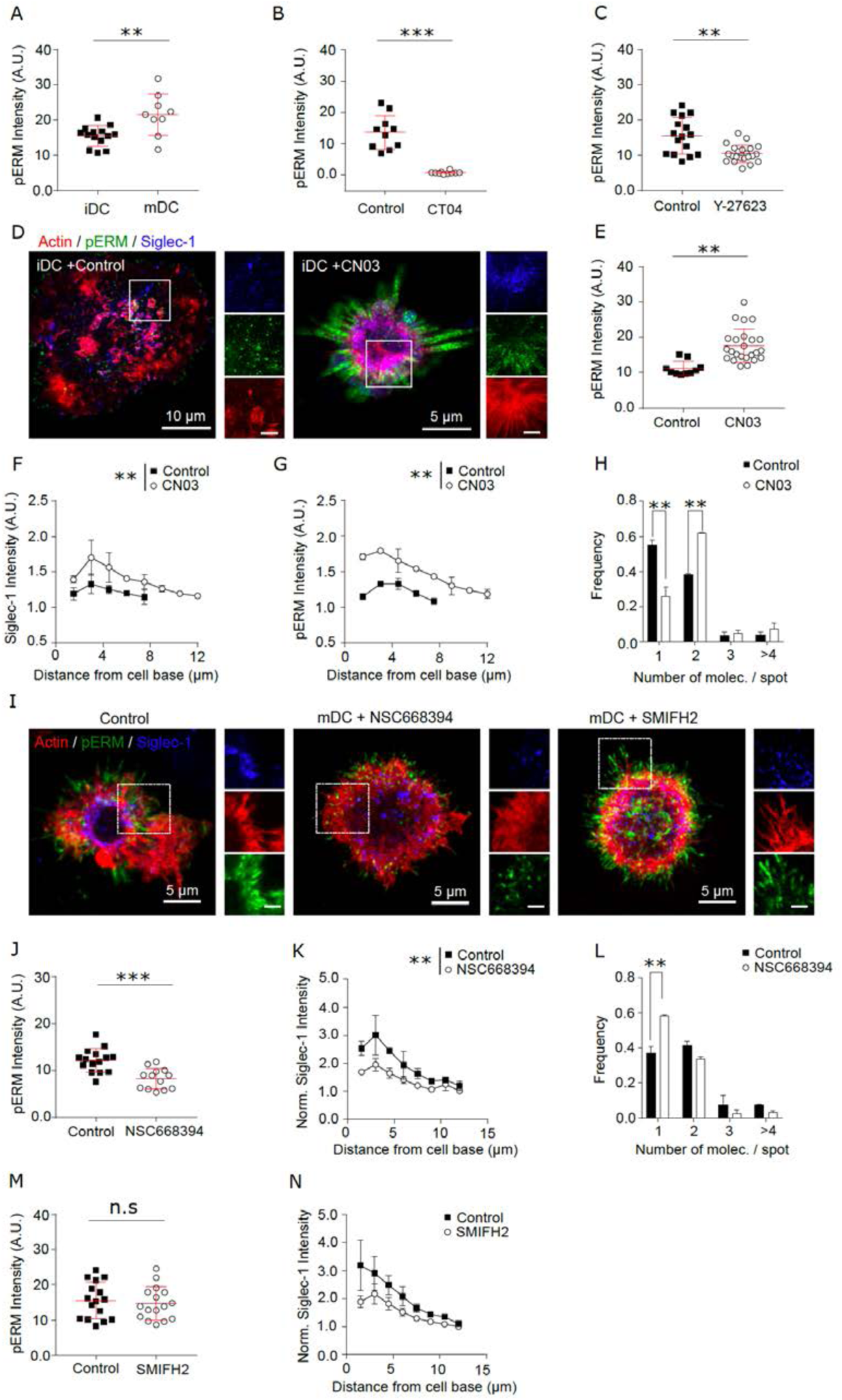
(A-C) pERM mean intensity values in control iDCs and control mDCs (A) or mDCs treated with CT04 (B) or Y-27623 (C). Results show the mean + S.D. of one representative experiment (N=2) in which a minimum of 9 (A), 7 (B) and 10 (C) cells per donor and condition were analyzed. (D) STED images of the basal plane for control iDCs or cells treated for 3h with CN04 (2 µg/mL) and immunostained with anti-Siglec-1, SiR-actin and pERM. Insets correspond to enlarged images of the regions highlighted in the main images (scale bars: 2 µm). (E) pERM mean intensity values in control iDCs and cells treated with CN03. Results show the mean + S.D. of one representative experiment (N=2) with a minimum of 9 cells/donor and condition. (F) Plots of the axial distribution of Siglec-1 intensity in control iDCs and cells treated with CN03. Values in each plane show the mean intensity in steps of 1.5 µm from the cell base, normalized by the minimum mean intensity within the whole stack. Each symbol represents the mean + S.E.M of two donors with at least 9 cells per condition. (G) Same as in (F) but for pERM intensity. (H) Frequency histogram of the number of Siglec-1 molecules/spot in control iDCs and cells treated with CN03. Mean + S.E.M of two donors, at least 9 cells/donor and condition. (I) Confocal maximum intensity projection images of control mDCs and cells treated for 3h with NSC668394 (250 µM), or for 1h with SMIFH2 (25 µM), immunostained with anti-Siglec-1, SiR-actin and pERM. Insets correspond to enlarged images of the regions highlighted in the main images. Scale bars: 2 µm. (J) pERM mean intensity values in control mDC and cells treated with NSC668394. Mean + S.D. of a minimum of 8 cells per condition from one representative experiment (N=2). (K) Same as in (F) but in control mDCs and cells treated with NSC668394. Each symbol represents the mean + S.E.M of two donors, with at least 9 cells per condition. (L) Frequency histogram of the number of Siglec-1 molecules/spot in control mDCs and cells treated with NSC668394. Mean + S.E.M of two donors, at least 9 cells/donor and condition. (M) pERM mean intensity values in control and SMIFH2 treated mDC. Data shows the mean + S.D. of one representative experiment (N=2) with a minimum of 10 cells per condition. (N) Same as in (K) but in control mDC and cells treated with SMIFH2. Each symbol represents the mean + S.E.M of two donors, with a minimum of 10 cells per condition.

**Figure supplement 4.**
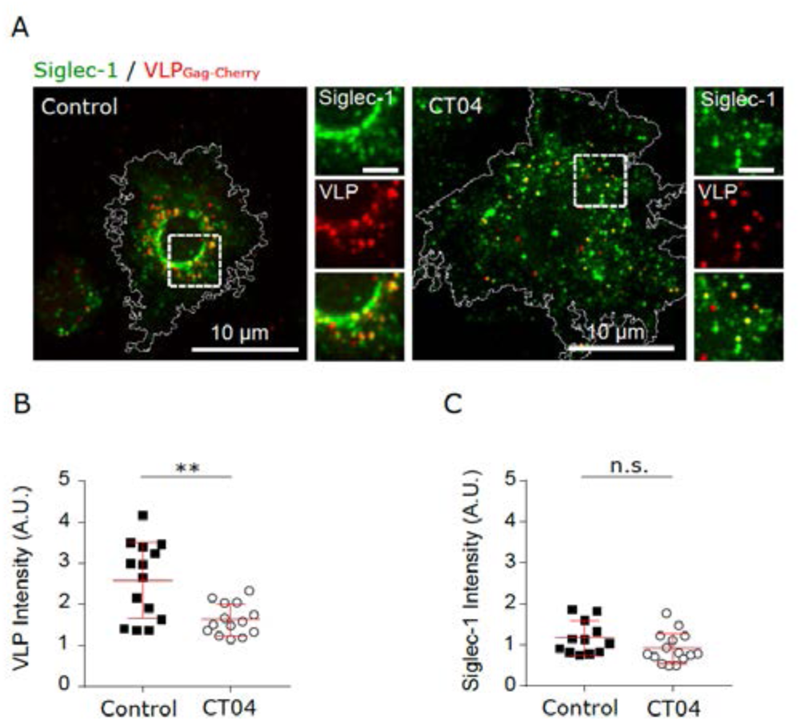
(A) Confocal images of control mDC, and cells treated with CT04, pulsed for 5 min with VLP-Gag-Cherry and immunostained with anti-Siglec-1. Insets on the right correspond to enlarged views of the regions highlighted in the main images (scale bar 2 µm). (B) VLP Intensity in control and CT04 treated cells, after 5 min of VLP capture. Mean + S.D. of one representative experiment (N=2) with 14 cells per condition. (C) Levels of Siglec-1 in control and CT04 treated cells, after 5 min of VLP capture. Mean + S.D. of one representative experiment (N=2) with 14 cells per condition.

**Figure supplement 5.**
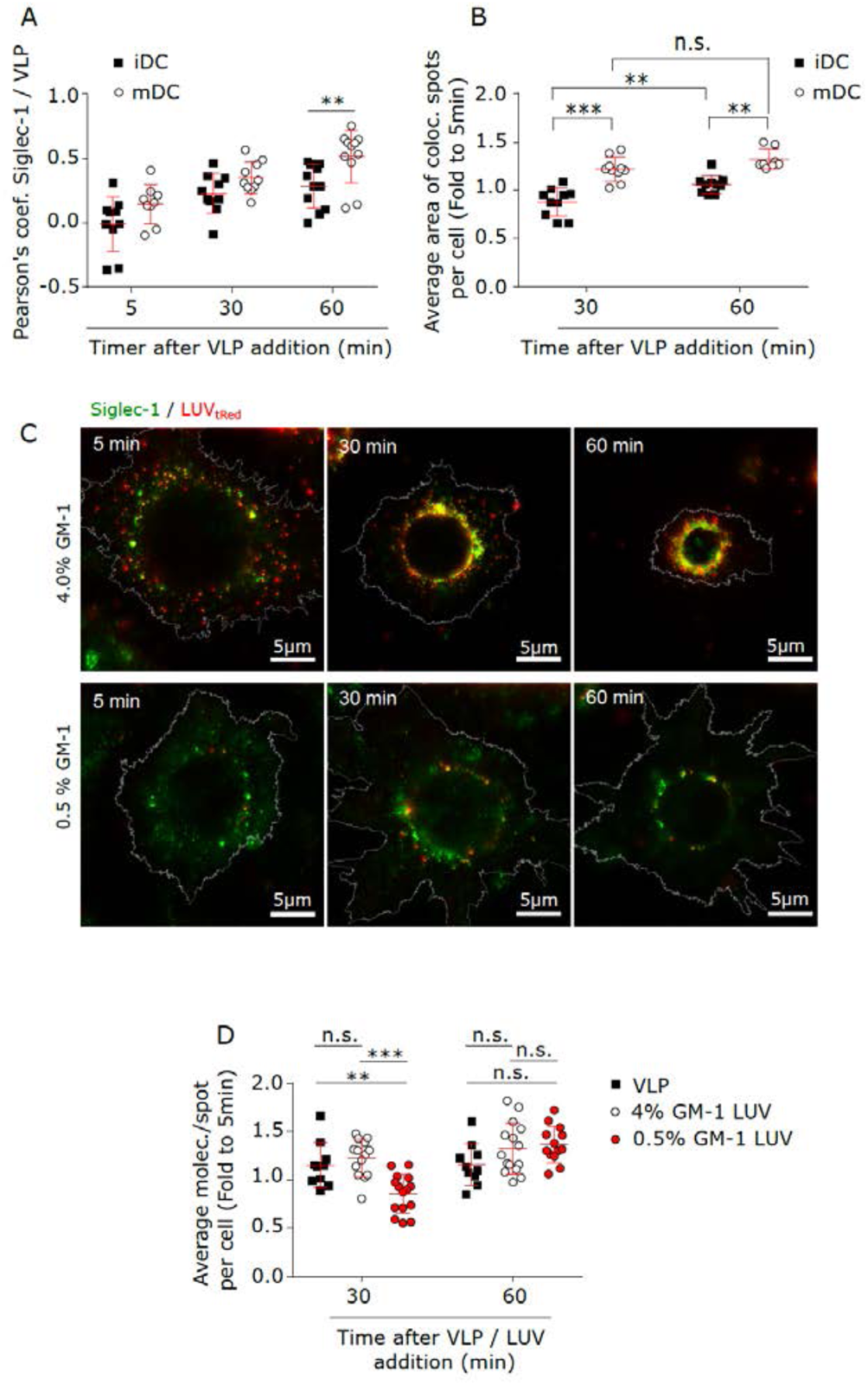
(A) Colocalization between Siglec-1 and VLP-Gag-Cherry in iDC and mDC after different times of VLP capture as determined by the Pearson coefficient. Mean + S.D. of a minimum of 9 cells per condition. (B) Size of Siglec-1 spots colocalizing with VLP-Gag-Cherry in iDC and mDC after different times of VLP capture. The average value of all colocalising spots in each cell after 30 and 60 minutes of VLP capture is expressed as a fold increase to the average value after a 5 min pulse. Mean + S.D. of a minimum of 8 per condition. (C) Representative STED images of mDC fixed and immunostained with anti-Siglec-1 after different times of incubation with LUVs carrying 4% (upper panels) and 0.5% (lower panels) GM-1. (D) Number of Siglec-1 molecules/spot not colocalizing with either VLP, 0.5% or 4% GM-1 LUVs in mDC. Each symbol represents the average value of all non-colocalizing spots in each cell after 30 and 60 min of VLP/LUV addition expressed as a fold increase to the average at 5 min. Mean + S.D. of a minimum of 10 cells per time and condition.

**Figure supplement 6.**
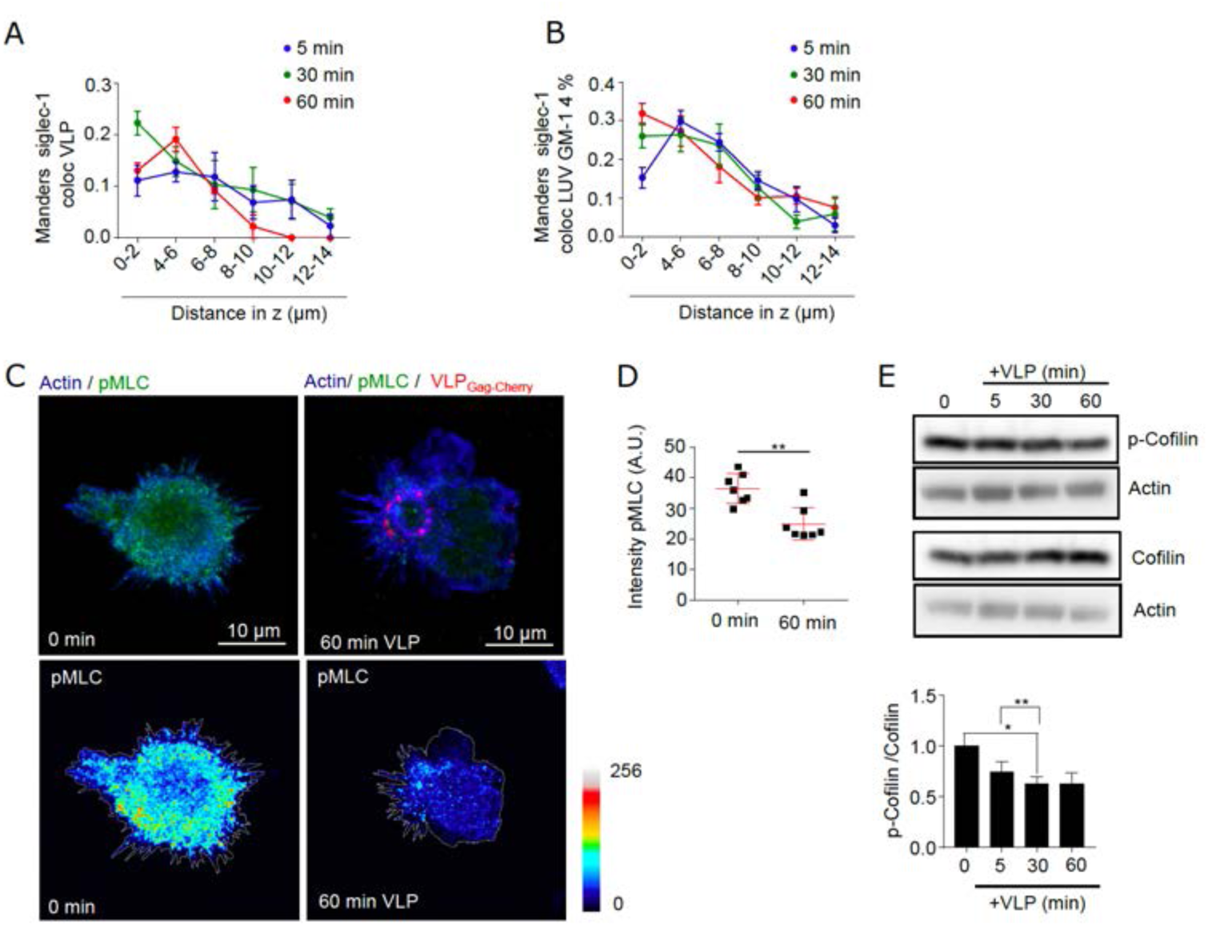
(A) Manders coefficients in the axial plane of Siglec-1 colocalizing with VLP-Gag-Cherry (representative images in Figure 6A). Data in (A) shows the mean + S.E.M of three (time points of 5 min and 30 min) and 2 donors (60 min) with a minimum of 9 cells per donor and time. (B) Same as in A but in cells incubated with 4.0% GM-1 LUVs. Data corresponds to the mean + S.D. of a minimum of 7 cells per time and condition. (C) Representative confocal images of mDC immunostained with anti-pMLC and SiR-actin at steady state and after 60 min of incubation with VLP-Gag-Cherry. Images in the bottom show projections of color-coded intensity values for pMLC. (D) pMLC intensity in mDC at steady-state and after 60 min incubation with VLP-Gag-Cherry. Data correspond to the mean + S.D. from one representative experiment (N=2) counting 7 cells per condition. (E) Representative immunoblots from cell extracts of mDC at steady-state and incubated in the presence of VLP-Gag-Cherry for different times. Equal amounts of protein loaded in each line were revealed with anti-cofilin, p-cofilin (S19) and b-actin. Bars in the bottom graph show the mean + S.E.M. of three independent experiments quantifying the ratio between p-cofilin and cofilin levels normalized to time 0 min.

